# Targeting of MMP-13 prevents aortic aneurysm formation in Marfan mice

**DOI:** 10.1101/2022.11.30.518511

**Authors:** Laura-Marie A. Zimmermann, Ariane G. Furlan, Dennis Mehrkens, Simon Geißen, Alexandra V. Zuk, Galyna Pryymachuk, Nadine Pykarek, Tim van Beers, Dagmar Sonntag-Bensch, Julia Marzi, Katja Schenke-Layland, Jürgen Brinckmann, Paola Zigrino, Maria Grandoch, Stephan Baldus, Gerhard Sengle

## Abstract

Fibrillin-1 assembles into microfibrils that not only define the structural integrity and biomechanics of the aorta but also target and sequester growth factors within the extracellular microenvironment of aortic resident cells. To better understand how dominant negative effects on fibrillin microfibril stability manifest in growth factor driven aortic disease, we analyzed early events of aortic aneurysm formation within the first two weeks of postnatal life in the dominant negative *Fbn1* GT8 Marfan mouse model. Echocardiography analysis of homozygous GT8 *Fbn1* mice showed significant aortic root enlargement within the second week of postnatal life which correlated with the onset of fibrillin-1 fiber degradation, aberrantly increased BMP activity and upregulated transcript levels of the collagenase MMP-13. We also found the aortic collagen network structurally disturbed where the mutant GT8-fibrillin-1 was detected. Genetic ablation or pharmacological inhibition of MMP-13 in *Fbn1* GT8 Marfan mice prevents aortic root dilatation implicating the relevance of this mechanism in aortic aneurysm formation in Marfan syndrome.

## Introduction

Supramolecular networks composed of fibrillins (fibrillin-1 and fibrillin-2) and associated ligands form intricate cellular microenvironments that balance tissue homoeostasis and direct remodeling (1). Fibrillins assemble into “beads-on-a-string” microfibrils that are not only indispensable for conferring elasticity to the aortic wall, but also control the bioavailability of growth factors targeted to the extracellular matrix (ECM) architecture (2). Fibrillin microfibrils (FMF) not only represent the structural core of elastic fibers, but also decorate the surface of elastic fibers and form independent networks. FMF have a diameter of 10-12 nm and show a bead-to-bead periodicity of about 50 nm which matches the collagen banding period (3). They exist in tissues as homo- and heteropolymers depending on the spatio-temporal context (4).

The importance of FMF for tissue homoeostasis is illustrated by the clinical features caused by mutations in the human fibrillin genes (*FBN1*, *FBN2*), summarized as “fibrillinopathies”. The fibrillinopathies represent disorders with distinctive, common, or opposing phenotypes affecting the musculoskeletal, cardiovascular, ocular, pulmonary, central nervous and dermal system. The most prevalent disorder caused by *FBN1* mutations is Marfan syndrome (MFS)(MIM#154700) which is characterized by long bone overgrowth (tall stature, arachnodactyly), lack of muscle tone, hyperelastic skin, and cardiovascular complications such as aneurysm formation at the aortic root and mitral valve prolapse. Until now, more than 3000 *FBN1* mutations in all exons have been identified leading to MFS while also rare *FBN1* mutations lead to long bone undergrowth (short stature, brachydactyly), hypermuscularity, thick skin, and no signs of cardiovascular involvement. This suggests that FMF control tissue homeostasis by balancing growth factor signals.

However, currently it is not well understood how specific structural alterations of the FMF ultrastructure lead either to aberrant growth factor activation or loss of their specific tissue targeting. Moreover, the structural integrity of FMF is also crucial for proper assembly and stability of other extracellular matrix (ECM) networks in connective tissues. Results from currently studied *Fbn1* mutant mouse models of MFS suggest that haploinsufficiency, the absence of one wild-type *Fbn1* allele, is the threshold requirement for balancing connective tissue homeostasis. This suggest that below 50% production of wild-type fibrillin-1 protein FMF growth factor sequestration is not guaranteed leading to their aberrant activating and respective detrimental downstream consequences. An alternative idea is that dominant negative effects on FMF assembly and stability are caused by the secretion of mutant fibrillin-1 that is incorporated into FMF and thereby compromises the structural integrity of the FMF ultrastructure.

To better understand how dominant negative effects on FMF stability manifest in growth factor driven aortic disease, we analyzed early events of aortic aneurysm formation within the first two weeks of postnatal life in the *Fbn1* GT8 Marfan mouse model. The *Fbn1* GT8 mouse model was generated by introducing a mutation in the “neonatal” region of fibrillin-1 leading to truncation of the C-terminal half. To follow the fate of the mutant fibrillin-1, an eGFP tag was placed at the end of the truncated molecule (5). It was shown that truncated GT8-fibrillin-1 is secreted at the same level as wild-type fibrillin-1 and is clearly assembled into microfibrils in tissues from *Fbn1* GT8 mice. FMF from homozygous GT8 mice showed a loss of their periodic labeling suggesting an abnormal integration of FMF into the surrounding collagenous environment (5).

Here, we shed new light on the functional interaction of the FMF and the collagen network in the aorta and proof mechanistically that collagenase antagonism represents a valid strategy to blunt aortic aneurysm growth in MFS.

## Materials and methods

### Mice

This study was carried out in strict accordance with German federal law on animal welfare, and the protocols were approved by the “Landesamt für Natur, Umwelt und Verbraucherschutz Nordrhein-Westfalen” (84-02.04.2014.A397 and 81-02.04.2019.A326 for breeding, 84-02.05.40.14.115 for euthanasia, and AZ: 84-02.04.2019.A033 for pharmacological treatment and echocardiography). All mice were kept in specific pathogen free (SPF) condition, light, temperature and humidity were monitored. GT8 and MMP-13 mice were maintained on a C57Bl6J background and genotyping by PCR was performed as described before (5, 6). For experimental purposes, young animals up to the age of 14 days were killed by decapitation and older animals by cervical dislocation according to the animal protection law.

### Antibodies

The following primary antibodies and dilutions were used for western blot (WB), immunohistochemistry (IHC), or immunofluorescence (IF) analysis: anti-Histone H3 (#9715, Cell Signaling, Danvers, MA, USA; WB: 1:500), anti-phopho-SMAD1 (#ab97689, Abcam, Cambridge, UK; IHC: 1.100), anti-phopho-SMAD5 (#ab92698, Abcam; IHC: 1.100), anti-phospho-SMAD1/5/8 (#9511, Cell Signaling, WB: 1:1000), anti-BMP-7 GF (#500-P198 B, Peprotech, Cranbury, NJ, USA, WB: 1:2000), anti-collagen I (#ab34710, Abcam, WB: 1:500; IF: 1:50), anti-collagen III (#1330-01, Southern Biotech, Birmingham, AL, USA; WB: 1:500, IF: 1:1000), anti-tropoelastin (#PR385, Elastin Products Company, Owensville, MO, USA; IF:1:200), anti-MMP-13 (#ab39012, Abcam, WB:1:500; IF: 1:250), anti-MMP13 (#262471, St John’s Laboratory, London, UK, WB: 1:1000), anti-GFP (#70R-GG001, Fitzgerald Industries International, Acton, MA, USA; IF: 1:400), and anti-decorin (#ENH019-FP (LF114), Kerafast, Boston, MA, USA; IF: 1:1000). Polyclonal rabbit anti-FBN1 antiserum (WB: 1:2000, IF: 1:1000) was raised against the recombinantly produced N-terminal half of human fibrillin-1 (F90) (7, 8). Anti-rabbit or –mouse IgG HRP-linked Antibody (#P0399 and #P0260, DAKO GmbH, Jena, Germany; WB: 1:400) and goat anti-Rabbit IgG Alexa Fluor 555 (#A-21428, Thermo Fisher Scientific, Waltham, MA, USA; IF: 1:400) were used as secondary antibodies.

### Mouse aorta preparation

After mice were sacrificed, the thorax was opened and the right ventricle of the heart was perfused with an insulin syringe coupled to a needle (4 mm) to wash the aorta with 1 × PBS. Afterwards, the right ventricle was either perfused with 1% agarose gel for subsequent immunofluorescence and immunohistochemistry analysis or aortas were snap-frozen in liquid nitrogen for RNA isolation and protein extraction. After agarose gel polymerization, the aorta was dissected from the aortic valve until the diaphragm.

### Sample dehydration and clearing

Dehydration and clearing of dissected mouse aortas were as previously described (9). After postfixation with 4% PFA/PBS for 2 hours, aortas were dehydrated with ethanol at pH 9.0 (4 hours in 50%, 4 hours in 70%, and 2 × 4 hours in 100%) at 4°C to 8°C in gently shaking 5-ml tubes. After dehydration, the samples were transferred to ethyl cinnamate (Eci) (#112372, Sigma-Aldrich, St. Louis, MO, USA) and incubated while gently shaking at RT until they became transparent.

### Paraffin embedding and histological staining

Dissected aortas were positioned in embedding chambers and fixed in 4 % PFA / PBS at 4 °C, overnight. Trichrome Stain (Masson) Kit (Sigma-Aldrich, Darmstadt, Germany) was used to detect collagens in blue on tissue sections. The staining procedure was carried out according to the manufacturer’s instructions in the following order: dewaxing, short washing step with distilled H_2_O, incubation with Bouin’s solution for 15 min at 55 °C, washing with tap water for 10 min, a short washing step with distilled H_2_O, 70 % EtOH / HCl (1 ml of 32 % HCl / 100 ml) 5 min, washing with running tap water for 5 min, with Biebrich Scarlet acid fuchsin solution for 5 min, distilled H_2_O for 5 min, phosphotungram-/ phosphomolybdic acid for 5 min, aniline blue for 5 min, 2 % acetic acid for 2 min, short washing step with distilled H_2_O, followed by a final dehydration step. Orcein staining was used to visualize elastic fibers in the ascending aorta of mice. Therefore, paraffin sections were incubated for 25 min in 1 % orcein staining solution at RT, followed by a short washing step, dehydration and capping.

### Immunofluorescence and immunohistochemistry

For immunofluorescence analysis of paraffin embedded aortas heat-induced epitope retrieval was performed. Therefore, PFA-fixed tissue preparations were subjected to epitope retrieval by boiling in an evaporation dish at 180 W for 15 min to avoid bubble formation with a citrate buffer at pH 6.0. The buffer level was checked every 5 min, and was refilled, whenever necessary. After boiling, the evaporation dish was cooled together with the slides for 20 min at RT. After a TBS washing step and a one-hour blocking step with 5 % BSA / 0.1% Triton X-100 in 1 × TBS, the sections were incubated at 4 °C overnight with the primary antibody solution and detected with secondary antibodies or with the chromogen AEC using the IHC Permanent AEC Kit (Zytomed Systems, Berlin, Germany). For immunofluorescence on cryosections, aortic or skin biposies were harvested and embedded in optimal temperature cutting (OCT) medium without fixation. For the detection of intracellular proteins, the tissue was permeabilized by adding Triton X-100 into the blocking solution (5 % BSA in PBS) for 15 min. Right after, a 1:1 mixture of ice-cold methanol and acetone was added for 10 min at -20°C or 4 % PFA in PBS at 37 °C for 10 min. After a short washing step with TBS, sections were edged with a liquid blocker pencil and dried for 5 min. The blocking step was carried out by incubation of 25-30 μl of blocking solution (5 % BSA in 1 × TBS) per section in a humid chamber for 1 h at RT. Subsequently, the primary antibody solution was applied to sections and incubated at 4 °C overnight. After three TBS washing steps for 10 min, the secondary antibody solution was added at appropriate dilutions and incubated at RT for 1 h. Finally, cell nuclei were counter-stained with DAPI and capping was performed in DAKO Mounting Medium. For immunofluorescence of collagen PFA fixed sections were subjected to epitope retrieval and subsequently treated with 1mg/ml pepsin at 37 °C for 15 min, 0.15 mg/ml hyaluronidase at 37°C for 10 min, and proteinase K (1:2000) at 55°C for 10 min.

### Second harmonic generation microscopy

For collagen detection, 10 µm crossections of paraffin embedded aortas were dewaxed, and second harmonic generation microscopy (SHG) of aortic collagens was obtained by using a Leica TCS SP8 microscopy (Leica microsystems, Wetzlar Germany) with a Chameleon Vision II laser. SHG and TPEF were excited at 810 nm and imaged with an IR Apo L25x/0.95 W objective lens. For eGFP excitation, a 488-nm optically pumped semiconductor laser was used.

### Electron microscopy analysis of aortic samples

For ultrastructural analyzes of elastic lamellae at P0 and P7 aortic samples were harvested and fixed in cacodylate buffer. Contrasting was carried out by addition of 1.5 % aqueous uranyl acetate solution for 15 min at 37°C followed by a washing step in distilled water for 4 minutes in lead nitrate. After a 2 hours incubation with 2% osmium (VIII) oxide in the dark at RT, dehydrogenation and embedding in propylene oxide took place. Ultra-thin sections with a thickness of 70 nm were produced with ultramicrotome (Leica EM UC6) and applied to carbon nanotubes coated with carbon-reinforced plastic film. EM images are taken with a JEM-2100 PLUS transmission electron microscope. For ultrastructural analysis of aortic walls at P9, mice were euthanized and quickly fixed in supine position. The abdominal was instantly opened, the diaphragm was incised and the thorax was opened. Left heart ventricle was cannulated with 26GA I.V. cannula (BD Vasculon™ Plus, ref 393300) followed by incision in the right atrium. Then the circulatory system was rinsed free of blood with saline solution that contained (in mmol/l) 5 D-glucose, 24 NaHCO_3_, 3,2 NaH_2_PO_4_·H_2_O, 115 NaCl, 1 MgCl_2_·6H2O and 4 KCl for 90 sec, and then perfused with 2,5% glutaraldehyde in 0.1M phosphate buffer for 5 min. Aortas were carefully dissected and postfixed by immersion in fresh 2% glutaraldehyde overnight at 4 °C. Then the liver specimens were thoroughly washed in cacodylate buffer and osmicated (1% OsO_4_ + 1.5% K_3_[Fe(CN)_6_]), dehydrated via ethanol and propylene oxide and embedded in epoxide resin (EmBed, Sigma Aldrich). The embedded aorta samples were used for performing of 0.5 μm semithin and 50 nm ultrathin sections with an ultramicrotome (Ultracut E, Reichert-Jung). The semithin sections were transferred to a drop of distilled water on a microscope slide (Epredia^TM^, SuperFrost^TM^ Plus, Thermofisher Scientific, Germany), stretched using chloroform vapour, dried on hot plate (80°C), stained with 1% toluidine blue/ 1% Borax solution (60 sec). The ultrathin sections were cut using a diamond knife (Diatome) and mounted on formvar-carbon-coated 200-mesh copper grids (Sigma-Aldrich, TEM-FCF200CU50) or on uncoated 200-mesh nickel grids (Sigma-Aldrich, TEM-FCF200NI). For staining, 2% uranyl acetate in 50% ethanol and 0.2% lead citrate pH 11.8 were used according to (10). Transmission electron microscopy was performed using the Zeiss electron microscope EM109 (80kV, 200 μm condenser and 30 μm objective apertures, 2k SSCCD camera (TRS)).

### Raman microspectroscopy

Raman measurements were performed with a confocal Raman microscope (WiTec alpha 300 R, WiTec GmbH, Ulm, Germany) equipped with a green laser (532 nm) as described previously (11, 12). Frozen cross sections of aortic tissues were rinsed with PBS and a 63x Apochromat water dipping objective (N.A. 1.0; Carl Zeiss GmbH) was utilized for data acquisition. High-resolution scans were performed on areas of 20 × 20 μm with 0.5 × 0.5 μm pixel resolution and 0.5 s acquisition time per spectrum. Laser power was set to 50 mW. Frozen sections were washed, and kept under PBS during the measurements. Spectral data were pre-processed by cosmic ray removal and background subtraction. The spectral maps were analyzed by True Component Analysis (TCA), a tool from Project FIVE 5.0 software (WiTec GmbH) that identified different spectral information within the dataset, as previously described (13). Raman spectral datasets were extracted for collagen fiber components identified by TCA. The data were normalized and the Raman shift range was cropped to the fingerprint region between 400 and 1800 cm^-1^. The Unscrambler X10.5 (CAMO Software, AS, Oslo, Norway) was applied to perform principal component analysis (PCA) using NIPALS algorithm.

### Collagen protein quantification and crosslinking analysis

For crosslink analysis, isolated aortic arches from P90 nice samples were reduced by sodium borohydride to stabilize acid-labile collagen crosslinks (Sigma-Aldrich; 25 mg NaBH_4_/ml in 0.05 M NaH_2_PO_4_/0.15 M NaCl pH 7.4, 1 hour on ice, 1.5 h at RT). Specimens were digested two times with “high purity bacterial collagenase” (Sigma-Aldrich; 50 U/ml, 37 °C, 18 hours). After centrifugation, the soluble fractions (collagen crosslinks) were hydrolyzed in 6 N HCl at 110 °C for 24 hours. The hydrolysates were pre-cleaned by solid phase extraction (Agilent, Santa-Clara, CA, USA). Dried eluates were dissolved in sodium citrate buffer (pH 2.2) and analyzed on an amino acid analyzer (Biochrom 30+, Biochrom, Cambridge, UK) in a three-buffer gradient method and post-column ninhydrin derivatization. Elution took place for 5 min (flow rate: 15 ml/hour) with sodium citrate buffer (pH 4.25), 40 min with sodium citrate buffer (pH 5.35) and 20 min with sodium citrate/borate buffer (pH 8.6) at 80 °C. Quantification was based on ninhydrin generated leucine equivalence factors (DHLNL, HLNL: 1.8; HP: 1.7; HHMD: 3.4)(14). The nomenclature used in this manuscript refers to the reduced variants of crosslinks (DHLNL: dihydroxylysinonorleucine, HLNL: hydroxylysinonorleucine, HHMD: histidinohydroxymerodesmosine). The determination of the amount of collagen and protein was carried out on the hydrolyzed samples before subjection to the solid phase extraction. The collagen content was calculated on the basis of 14 mg hydroxyproline per 100 mg collagen.

### Quantitative real-time PCR

Isolation of RNA from primary murine VSMCs or isolated aortic arch at P14 was carried out by applying the phenol-chloroform extraction method using the TRIzol^TM^ reagent (Thermo Fisher Scientific, Waltham, MA, USA) according to the manufacturer’s instructions. The isolated RNA was air-dried after precipitation and taken up in 20 μl of RNase-free water followed by a subsequent sample purification using the RNeasy kit (Qiagen, Venlo, The Netherlands). Residual DNA contamination was removed from each sample using the Turbo DNA-free kit (Ambion, Austin, TX). RNA was quantified by photospectrometry, and 1.0 μg of RNA per sample was reverse-transcribed was reverse transcribed using the iScript™ cDNA synthesis kit from Bio-Rad (Bio-Rad, Hercules, CA). Quantitative PCR was performed using SensiFAST SYBR Hi-ROX Kit in 25 µL reaction volume (Meridian Bioscience, Cincinnati, OH). PCR was conducted with the StepOnePlus system (Applied Biosystems, Thermo Fisher Scientific). The standard annealing temperature of 60°C was chosen. Analysis of data was performed using the 2^−ΔΔCt^ method (15) and quantitated relative to the murine *Arbp* gene.

### VSMC isolation and stimulation

Primary VSMC were isolated from the aortic arches of P14 mice. After isolation, aortic arches were cut lengthwise to expose the intimal layer and immersed in collagenase solution (300 U/ml of collagenase in HBSS buffer) for 10 min at 37 °C. The adventitial layer was removed with a very thin tweezer. Aorta was cross-cut in small pieces and placed in 12-well plates in DMEM containing 20 % FCS at 37 °C overnight. Next day, aortic pieces were transferred to a collagenase-elastase solution (200 U/ml collagenase, 0.4 U/ml elastase in HBSS medium) for 45 min at 37 °C. Afterwards, aortic pieces were squeezed by repetitive pipetting using 1 ml tips in a first step and 200 µl tips in a second step. The reaction was stopped by adding 3 ml/well of DMEM containing 20 % FCS and the wells were incubated in this medium for 48 hours. As soon as cells achieved confluence on 12-well plates, they were transferred to a 25 mm^2^ flask and were used for experiments from the third passage on. For BMP stimulation, 1×10^5^ VSMC were seeded onto 6-well plates and cultivated for 4 days. On the fourth day, cells were stimulated in serum free media containing 100 ng/ml BMP-7 GF or 0.1 % BSA as negative control for 24 hours. After stimulation, cells were washed with PBS followed by RNA isolated using the TRIzol^TM^ reagent.

### Immunofluorescence analysis of ECM assembled in cell culture

5 × 10^5^ primary cells isolated from GT8 mice were seeded to 12 mm round coverslips in a 24-well plate. After four days of culture, cells attached to coverslips were washed with 1 × PBS and subsequently fixed with either a 1:1 mixture of ice-cold methanol and acetone for 10 min at -20°C or with 4 % PFA in PBS at 37 °C for 10 min. Coverslips were transferred to a humid chamber and blocked with 25 μl of 5 % BSA in 1 × PBS for 1h at RT to avoid non-specific binding. Subsequently, primary antibodies (table 3.5) were diluted in 1 % BSA and 25 μl of this solution was applied to the coverslips overnight at 4 °C. Secondary antibodies and the double-stranded DNA-binding core dye DAPI (4,6-diamidine-2-phenylindole dihydrochloride) were diluted in 1 % BSA and added to the coverslips for 1 h in the dark. After fixation, coverslips were washed three times with PBS before primary and secondary antibody incubations. Finally, samples were capped in DAKO Mounting Medium and analyzed by a fluorescence microscope (Axiophot, Zeiss) at 40 × magnification or a confocal laser scanning microscope (SP5, Leica).

### Co-culture assay

Primary dermal fibroblasts were isolated from newborn mice as previously described (16). Cells were cultured in Dulbecco’s Modified Eagle’s medium (DMEM GlutaMAX, Invitrogen, Carlsbad, CA) supplemented with 1% penicillin-streptomycin and 10% fetal bovine serum. For fibrillin-1 fiber formation cells were plated at a concentration of 1 × 10^5^ cells per well and grown on uncoated glass coverslips in a 24-well plate for 4 days. For co-culture assays (at 1:1 ratio), HEK-293-EBNA cells overexpressing BMP-7 complex (1 × 10^5^ cells per well) and wild-type or GT8 dermal fibroblasts (1 × 10^5^ cells per well) were plated together on uncoated glass coverslips in a 24-well plate. Slides were fixed in cold methanol, acetone (1:1) for 10 minutes at -20°C and non-specific binding sites blocked with 1% BSA/ PBS for 10 min RT. The cell layers were incubated for 1 hour with 10 µg per ml monoclonal anti-His6-tag antibody (R&D Systems) or a 1:1000 dilution of pAb 9543 279 anti-fibrillin-1. Slides were washed in PBS, and then incubated for 1 hour with Alexa Fluor 488 280 goat anti-mouse IgG and/or Alexa Fluor 555 donkey anti rabbit IgG at 1:1000 in 1% BSA/ 281 PBS. Slides were coverslipped using the Dako Fluorescence Mounting Medium (Dako, Denmark) and viewed using a Zeiss Axiophot microscope (Zeiss, Germany).

### Skin organotypic 3D co-culture

The employed 3D co-culture system was previously described (17). In brief, mouse wild-type keratinocytes (1 × 10^6^ cells/ml) were grown submersed on collagen I gels populated with mouse skin fibroblasts (5 ×10^5^ cells/ml) from *Fbn1^+/GT8^*, and *Fbn1^GT8/GT8^*mice, using 25-mm filter inserts. Fibroblasts were prepared from newborn mouse skin as previously described (16). The co-cultures were placed in 6-deep well plates. After 1 day, they were exposed to the air-liquid interface and medium change was performed every second day. The cell culture media after 21 days of culture was applied to seeded C2C12 cells for 5 hours followed by mRNA isolation and qPCR of *Id3* expression as previously described (18).

### Western blot and dot blot analysis

Aortic or skin biopsies and cell layers were incubated with lysis buffer (10 mM NaCl, 1.5 mM MgCl2, 20 mM Hepes pH 7.4, 20% glycerol, 0.1% Triton X 100 and 1 mM DTT) supplemented with 1× PhosSTOP (Roche, Switzerland), centrifuged at 440 × g for 5 min at 4° C, and washed with PBS before subjection to SDS-PAGE followed by western blotting using either a 1:1 mixture of rabbit pAb anti-SMAD1 (#ab63439, Abcam) and rabbit mAb anti-pSMAD5 (#ab92698, Abcam) or rabbit pAb anti-phospho SMAD1/5/8 (#9511, Cell Signaling) at 1:1000. For dot blot analysis of BMP-7 in conditioned media from co-culture assays 5 µl were applied to a nitrocellulose membrane and BMP-7 was detected using a rabbit pAb anti-BMP-7 GF at 1:2000 (#500-P198 B, Peprotech). For dot blot analysis in conditioned media of 3D organotypic co-cultures, 10 ml of media was concentrated over 2 ml heparin affinity medium (Heparin Sepharose 6 Fast Flow, GE Healthcare) packed into a 10 ml Poly-Prep Chromatography gravity flow column (Bio-Rad). After several washes with PBS, BMP-7 was eluted with 2 M NaCl.

### Pharmaceutical inhibitor treatment

The MMP-13 antagonist (BI-4394) has a very high potency to inhibit MMP-13 activity (IC_50_ = 1nM) (19). BI-4394 and the negative compound BI-4395 were applied to *Fbn1^+/GT8^*mice by osmotic minipumps (ALZET osmotic minipumps, Model 1004, 100 L) at concentrations of 1 mg/kg/day. Therefore, both compounds were dissolved in DMSO (1%) and diluted into sterile PEG 400 to a total volume of 100 _µ_L prior to filling into minipumps with a syringe under sterile conditions. Before implantation, minipumps were incubated in sterile PBS (25 mL) for 48 hours at 36 °C. Earliest implantation time point in mice was P30 and inhibitor/compound concentration was adapted to bodyweight (1. implantation bodyweight of: 1 18g, 2. 25g, 3. 30g). Ultrasound measurement of heart and aorta were performed as baseline before implantation. A subcutaneous pocket was performed in the neck of the animal and the minipump was implanted on the flank of the animal. Post operation pain killer were applied for 48 hours through drinking water (Buprenorphine 0.3 mg/ml, Temgesic). Minipumps released 0.11 L/hour of inhibitor or negative compound and stayed for 28 days in each mouse until reimplantation.

### Echocardiography

In order to measure the aortic diameter of mice animals were anaesthetized with isoflurane via mask ventilation and placed on a heating pad at 37°C. The abdominal skin was depilated and an ultrasound gel was applied on the naked abdomen. Subsequently, the ascending aorta was monitored using an ultrahigh frequency imaging platforms (VisualSonics Vevo 2100). B-mode recordings were performed using a MS 400 transducer (18–38 MHz). All images were recorded digitally and analysis was performed using the Vevo 2100-software. Calculations were performed with SPSS version 23.0, one way ANOVA and Turkey’s multiple comparisons test. After the recordings, mice were sacrificed and the aorta was harvested for immunofluorescence analysis.

### Statistical analysis

Data were represented as mean ± SD. Statistical analysis was performed using Prism 9 software (GraphPad, La Jolla, CA, USA). If the data followed normal distribution, statistical significance was determined by unpaired or paired t test for two-group comparisons and one-way/two-way analysis of variance (ANOVA) for comparison among three or more groups followed by Bonferroni’s correction for multiple comparison tests. If the normality assumption was violated, nonparametric tests (Mann-Whitney or Kruskal-Wallis) were conducted. Data were considered statistically significant for P values of 0.05 or less.

## Results

### Initiation of degradative processes on fibrillin-1 network in *Fbn1* GT8 aorta at P7

It was described that despite the presence of two truncated alleles *Fbn1^GT8/GT8^* mice showed a normal FMF network at birth (5). This suggests that during development and until birth fibrillin-2 is structurally and functionally compensating for the lack of full length fibrillin-1. Since the assembly process of fibrillin-1 monomers into microfibrils involves C- and N-terminal head-to-tail interactions (20) between monomers, truncation of the fibrillin-1 C-terminal half in GT8 mice affects FMF integrity in homozygotes at the time when fibrillin-2 production ceases. We therefore wanted to investigate at which postnatal time point degradative processes of FMF are initiated. Immunohistochemistry analysis of paraffin-embedded aortic cross sections using a polyclonal antibody against-fibrillin-1 showed that fibrillin-1 fibers were present in all aortic layers at birth (postnatal day: P0) and P7, however at P14 fibrillin-1 signals could only be detected in the adventitia. This might be due to reduced fibrillin-1 epitope availability caused by tropoelastin deposition onto FMF followed by crosslinking giving rise to mature elastic lamellae in the aortic wall. In *Fbn1^GT8/GT8^* mice, fibrillin-1 fibers appeared to be normal until P7, but their presence was strongly diminished at P14 (Fig. 1A).

**Figure 1:**
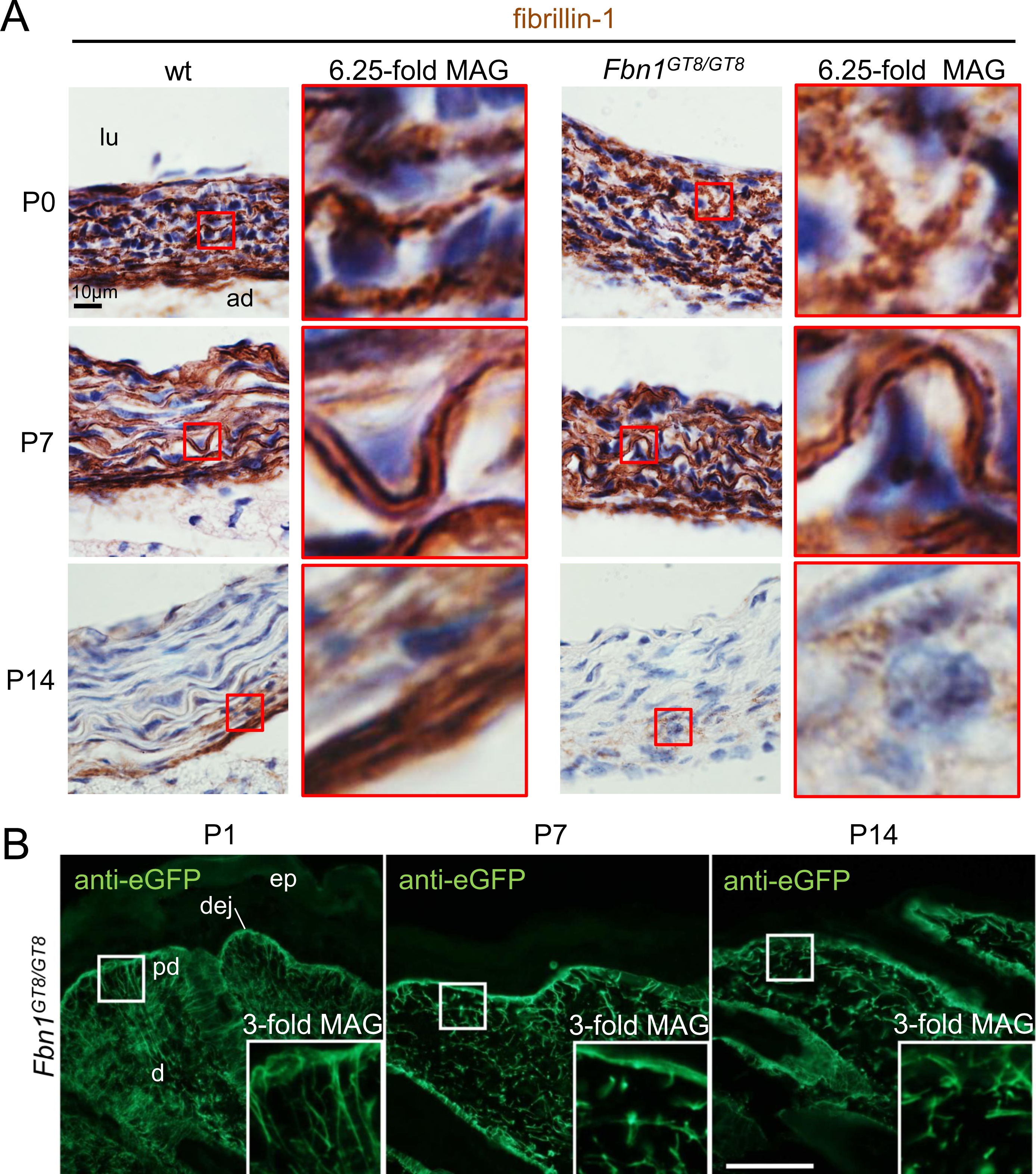
Initiation of degradative processes on fibrillin-1 network in *Fbn1* GT8 aorta and skin. (A) Fibrillin-1 expression in ascending aorta was monitored at P0, P7, and P14. Immunohistochemistry using a specific polyclonal antibody against-fibrillin-1 showed localization of fibrillin-1 throughout the aortic wall of wild-type (wt) mice at P0 and P7, however at P14, fibrillin-1 was only detectable in the adventitia. At P7, fibrillin-1 signals in wt and *Fbn1^GT8/GT8^* aortic walls were similar. At P14, fibrillin-1 signals in *Fbn1^GT8/GT8^* aortas was strongly reduced. lu: lumen, ad: adventitia. Scale bar: 10 µm. (B) eGFP positive fibrillin-1 fibers in skin from *Fbn1* GT8 mice are proteolytically degraded at P7, and P14 as shown by immunofluorescence (d: dermis, dej: dermal-epidermal junction, ep: epidermis, pd: papillary dermis). Inset: Boxed area is shown at 3-fold magnification. Scale bar: 50 µm.

Since FMF networks independent of elastin are difficult to analyze in the aorta we investigated fibrillin-rich oxytalan fibers that transverse from the dermis via the papillary dermis and insert perpendicular at the dermal-epidermal junction (dej) into the basement membrane. As eGFP fluorescence was shown to be only sufficient to detect signals at one month of age in GT8 tissues (5), we performed immunofluorescence staining using an anti-eGFP antibody. Our analysis of *Fbn1^GT8/GT8^* skin revealed that at P0 only intact eGFP-positive fibers could be observed. However, at P7 and P14 we observed a fragmented and punctate staining pattern confirming that degradative processes on FMF in tissues of *Fbn1^GT8/GT8^* are initiated in the second week of postnatal life (Fig. 1B).

Since fragmentation of elastic fibers after two months has been documented in *Fbn1^+/GT8^* mice (5) as well as in other *Fbn1* mouse models (21, 22), the status of elastic lamellae in the aortic wall of *Fbn1^GT8/GT8^* mice was investigated in the early postnatal period. To determine the earliest time points of visible elastic fiber degradation, elastic lamellae in P7 thoracic aortic walls were analyzed by light microscopy after Orcein staining of paraffin-embedded aortic sections. Elastic fiber breaks could be detected already at P7 in *Fbn1^GT8/GT8^* aortic walls and were more prominent at P14 (Fig. 2A).

**Figure 2:**
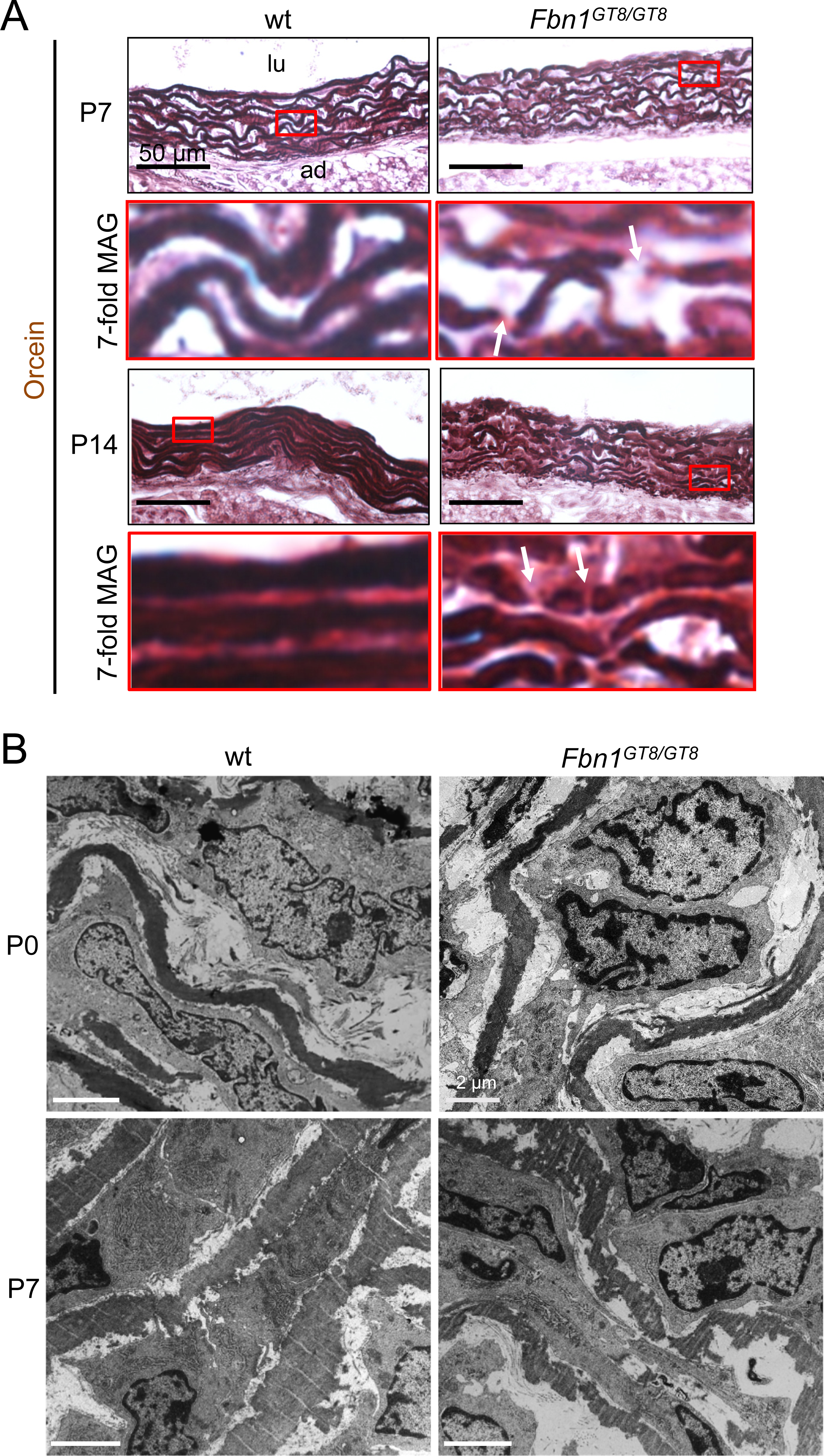
Elastic fiber breaks in *Fbn1* GT8 aortas are firstly detected at P7. (A) Orcein staining shows breaks in elastic fibers of *Fbn1^GT8/GT8^* at P7, which were more prominent at P14. Arrows indicate elastic breaks. Scale bar: 50 µm. (B) Transmission electron microscopy of GT8 perinatal aorta shows elastic fiber breaks at P7. Elastic fiber ultrastructure in *Fbn1^GT8/GT8^* aortas at P0 do not display any alterations. At P7 elastic fibers in *Fbn1^GT8/GT8^* aortic walls appear fragmented and disconnected to VSMCs. Polymerized, amorphous elastin was visualized in black after staining with osmium tetroxide.

Previous reports showed that elastic fibers in ascending mouse aorta can be detected by electron microscopy as early as E18, whereby the peak of elastin expression is achieved at P14 before it subsequently decreases (23). Transmission electron microscopy (TEM) analysis of *Fbn1^GT8/GT8^* aortic walls did not detect any ultrastructural alterations in elastic fiber integrity at P0, suggesting that absence of the C-terminal half of fibrillin-1 does not affect elastogenesis in comparison to mice deficient in FMF associated molecules such as fibulin-4, -5, EMILIN-1, or LTBP-4 (24–29). However, at P7 elastic fiber disruptions and disconnections between elastic fibers and VSMCs were already visible (Fig. 2B). In addition, TEM analysis of P9 aortic sections revealed discontinuous elastic lamellae and increased spacing of the intra lamellar space in *Fbn1^GT8/GT8^* aortas leading to a reduced physical contact of VSMCs with adjacent elastic lamellae. Closer inspection did not reveal any major morphological changes between VSMCs in *Fbn1^GT8/GT8^* and wild-type control aortas (Fig. S1).

### Diminished collagen in aorta of *Fbn1^GT8/GT8^* mice

It is known that fibrillin fibers together with elastic fibers provide resilience to the walls of arteries, whereas deposition of collagens fibers between the elastic lamellae as well as in the adventitial layer is required to confer tensile strength to the aorta (30). Since profound defects in the collagen microarchitecture have been found in thoracic aortic aneurysms of MFS patients (31), we analyzed collagen deposition in aortas of *Fbn1^GT8/GT8^* mice. Histological analysis using Massońs trichrome stain on paraffin sections of thoracic aortas showed decreased collagen deposition in the *Fbn1^GT8/GT8^* aorta at P7 and P14 (Fig. 3A). Second harmonic generation (SHG) microscopy utilizing two-photon excitation imaging to visualize the collagen fiber network confirmed that the amount of collagen in the adventitia of *Fbn1^GT8/GT8^* mice was significant reduced (Fig. 3B). In addition, collagen fibers in *Fbn1^GT8/GT8^* adventitia appeared not properly deposited since in contrast to the wild-type littermates no co-localization between collagen and elastin immunofluorescence signals was detected employing an antibody against tropoelastin (Fig. 3).

**Figure 3:**
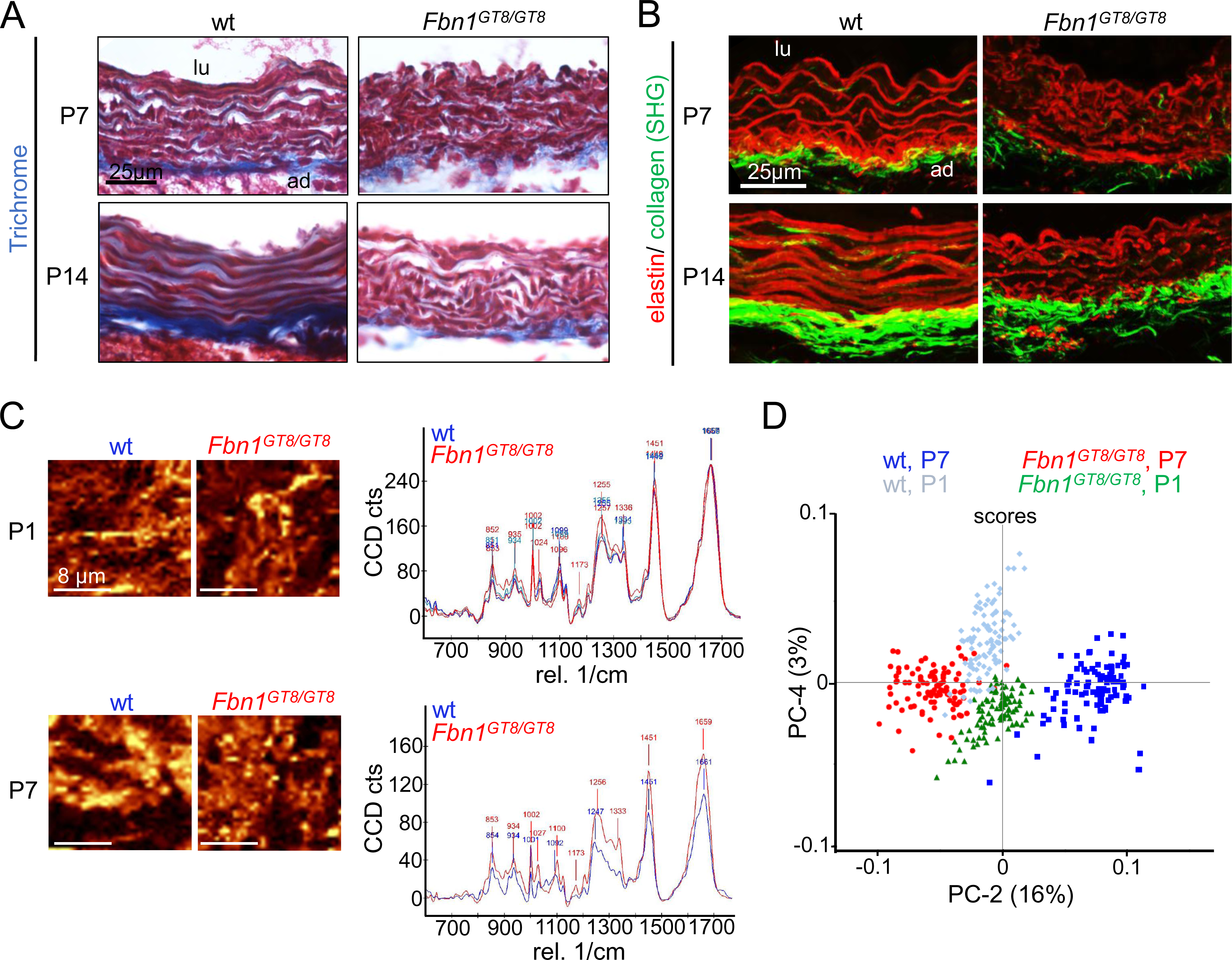
Trichrome stain, second harmonic generation microscopcopy, and Raman microspectroscopy of *Fbn1^GT8/^*^GT8^ aortas reveals significant change in adventitial collagen network at P7. (A) Histological analysis using Massońs trichrome stain on paraffin sections of thoracic aortas showed decreased collagen deposition in *Fbn1^GT8/GT8^* aorta at P7 and P14. Wild-type (wt) mice at P14 showed a striking gain in deposited collagen in the adventitia when compared to P7. A similar gain in adventitial collagen was not observed in *Fbn1^GT8/GT8^*aortas. (B) Paraffin embedded sections of thoracic aortas were harvested at P7 and P14 followed by immunofluorescence analysis with an antibody against tropoelastin. Simultaneously, second harmonic generation (SHG) microscopy utilizing two-photon excitation imaging was employed to visualize the collagen fiber network. At both time points, *Fbn1^GT8/GT8^*aortic walls showed increased elastic fibers breaks and strongly reduced collagen deposition when compared to wt controls. Moreover, co-localization between collagen and elastin signals found in adventitia of wt mice at P7 and P14 was abolished in *Fbn1^GT8/GT8^* mice. (C) (left) Visualization of Raman signals derived from representative individual collagen fibers in aortic adventitia measured on sections of aortic arches. Scale bars: 8 µm. (right) Overlay of Raman spectra of adventitial collagen fibers does not show any changes of molecular signatures at P1 but at P7. (G) Principal component analysis (PCA) of Raman data sets from wt and *Fbn1^GT8/GT8^* aortic adventitia shows a statistically significant difference at P7 but not at P1. Three mice per genotype were analyzed.

To further investigate at what time point collagen alterations occur in *Fbn1^GT8/GT8^* mice, we utilized Raman microspectroscopy on frozen sections of aortas from P0 and P7 mice. Raman microspectroscopy relies on the inelastic scattering of laser light and is capable to monitor changes of molecular signatures in cells and tissues before they can be detected by standard microscopy Lit. Our Raman analysis of collagen fibers in the adventitial layer of *Fbn1^GT8/GT8^* aortas visualized by SHG showed that at P0 no changes of collagen specific spectra was visible as indicated by principal component analysis (PCA) (Fig. 3C), while a statistically significant difference was observed at P7 (Fig. 3D).

### Fibrillin-1 shows co-distribution with aortic collagen which is diminished where mutant fibrillin-1 is present

Similar to our findings in GT8 aortas, a dramatically disturbed collagen architecture in the adventitial layer was also found in the aortic wall of MFS patients (31). This suggests that a preserved FMF integrity is required for proper collagen network formation in the adventitia. We therefore investigated the co-distribution of collagen and fibrillin-1 in the ascending aorta. By using optical clearing upon ethyl cinnamate treatment of dissected aortic arches followed by label free two-photon excitation imaging we were able to visualize collagen via the SHG signal and GT8-fibrillin-1 via the eGFP signal (Fig. 4A). This analysis revealed diminished collagen signals in the aortic root and ascending aorta, where also the presence of the mutant GT8-fibrillin-1 was detected. To investigate whether fibrillin-1 shows a potential co-localization with collagen networks in the aorta we attempted double immunolabeling monitored by confocal immunofluorescence microscopy (Fig. 4B). Since fibrillin-1 immunosignals are very difficult to detect in the aortic wall already at P14 (Fig. 1), we optimized epitope retrieval methods by including enzymatic digests of pepsin, hyaluronidase, and proteinase K subsequent to boiling aortic sections in citrate buffer. Thereby, we found that only aortic sections from aged mice show sufficient stability to obtain interpretable signals. Immunofluorescence analysis of *Fbn1^+/GT8^* aortas at one year of age showed that the content of collagen III in the aortic adventitia is significant diminished (Fig. 4C). Moreover, when performing co-localization analyses we found that fibrillin-1 shows the strongest co-localization with collagen III in the adventitial layer while co-localization in the medial layer only occurred partially (Fig. 4D).

**Figure 4:**
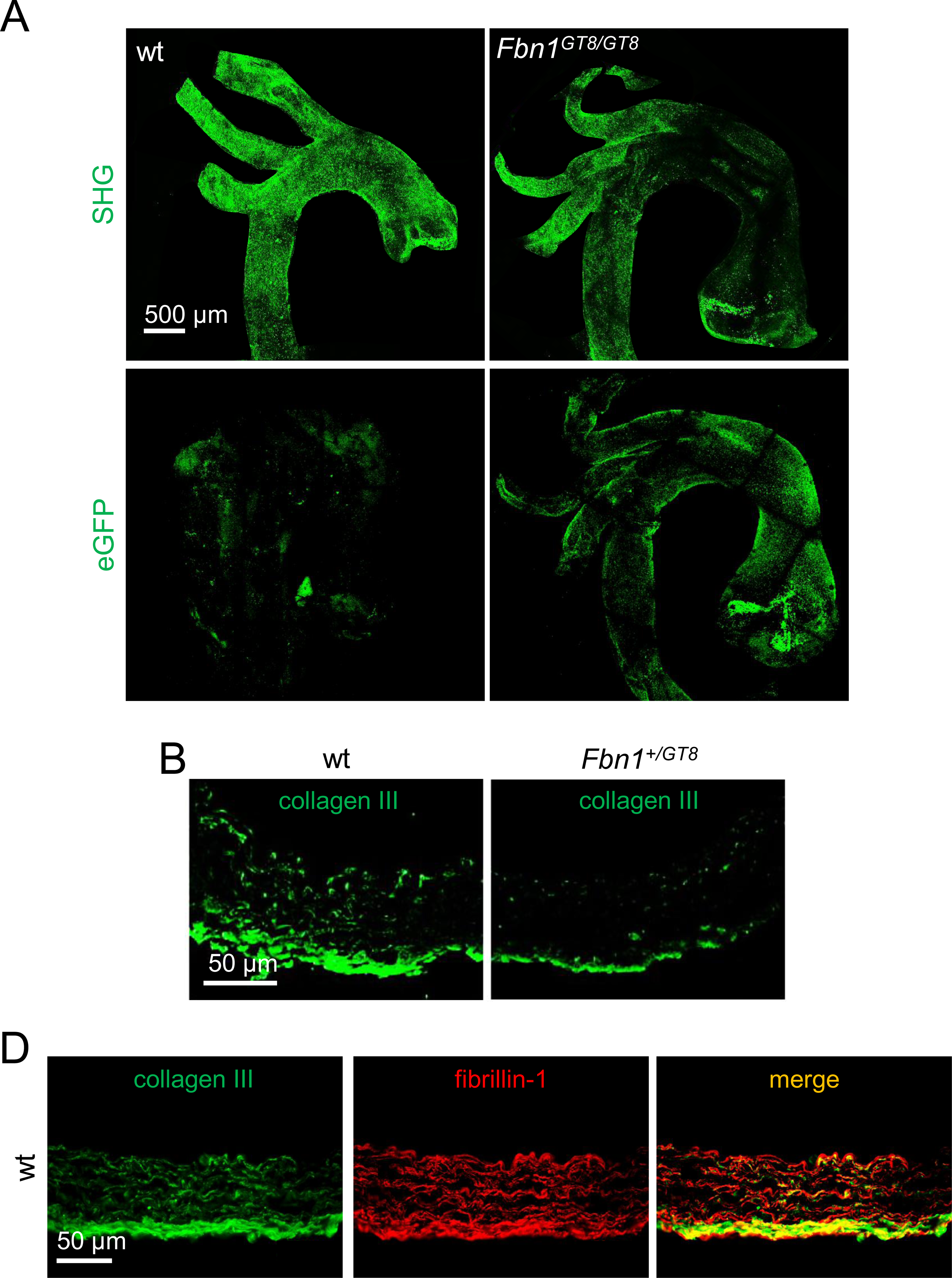
Fibrillin-1 shows co-distribution with aortic collagen which is diminished where mutant fibrillin-1 is present. (A) 3D imaging of whole P11 aortas after clearing with ethyl cinnamate (Eci) reveals reduction of collagen second harmonic generation (SHG) signal in the ascending aorta of *Fbn1^GT8/GT8^*mice where mutant e-GFP labelled GT8-fibrillin-1 was detected. Scale bar: 500 µm. (B) Confocal immunofluorescence microscopy show reduction of collagen III signals in aortic media and adventitia of *Fbn1^+/GT8^* mice at one year of age. Scale bar: 50 µm. (C) Confocal immunofluorescence microscopy after epitope retrieval on aortic sections of one year old wt mice indicates strong co-distribution of fibrillin-1 with collagen III in adventitia and partial co-localization in aortic media. Scale bar: 50 µm.

### Structural integrity of fibrillin-1 fibers is required to control BMP activity

Our results and previously obtained data from *Fbn1* GT8 mice (5) suggest that as long as FMF integrity is preserved by structural compensation via fibrillin-2 during the first week of postnatal life, elastic and collagen fibers do not show any structural alteration. Since in addition to their tissue specific architectural function, FMF also integrate ECM-cell communication by targeting growth factors of the TGF-β superfamily we next investigated whether the observed structural changes of fibrillin fibers correlated with the onset of dysregulated TGF-β or BMP activity. Previous studies showed that haploinsufficiency of fibrillin-1 leads to aberrant activation of TGF-β in the aorta at 14 weeks of age (21). In addition, our previous work demonstrated that FMF also directly target and sequester bone morphogenetic protein (BMP) prodomain-growth factor complexes (PD-GF CPLXs) (7, 32, 33). This implies that the presence of GT8-fibrillin-1 will structurally compromise FMF leading to failed sequestration of BMP PD-GF CPLXs. If binding to fibrillin-1 fibers confers latency, disruption of fibrillin-1 fibers will lead to the presence of unbound bioactive BMP complexes.

In order to address whether disturbed BMP sequestration is caused by the presence of mutant GT8-fibrillin-1 we analyzed cell cultures of primary cells from *Fbn1* GT8 mice. Co-cultures of wild-type murine dermal fibroblasts and BMP-7 CPLX expressing HEK-293-EBNA cells, which do not assemble ECM fibers, showed BMP-7 CPLX deposition on newly formed fibrillin-1 fibers (Fig. 5A). Immunofluorescence analysis of *Fbn1* GT8 fibroblast cultures showed that GT8-fibrillin-1 interferes with the assembly of wild-type fibrillin-1 in *Fbn1^+/GT8^*fibroblast cultures, while truncated GT8-fibrillin-1 cannot be assembled into fibrillin fibers in *Fbn1^GT8/GT8^* cultures (Fig. 5B).

**Figure 5:**
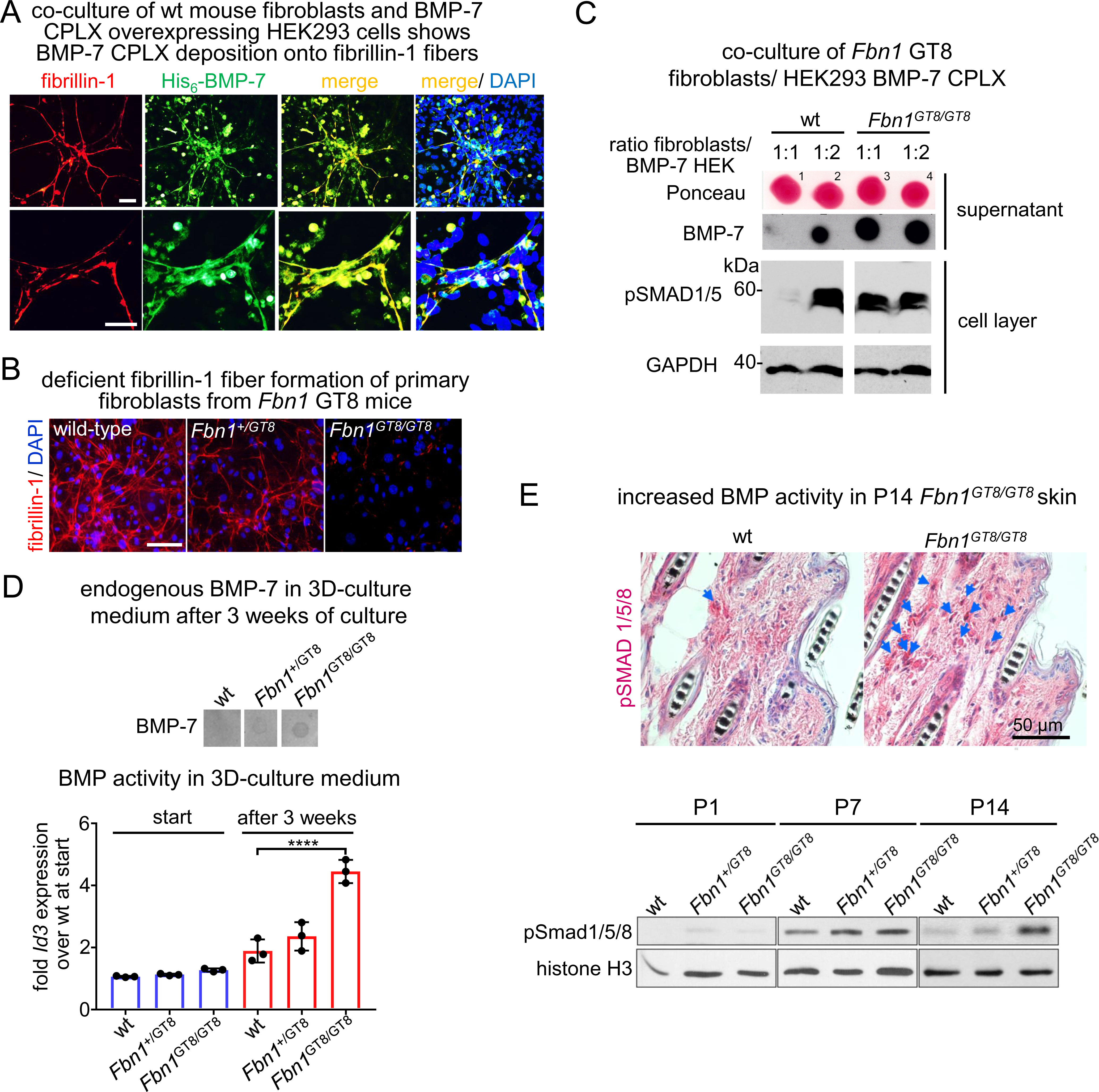
Structural integrity of fibrillin-1 fibers is required to control BMP activity. (A) Immunofluorescence analysis of co-cultures of wild-type (wt) murine primary skin fibroblasts and His_6_-tagged BMP-7 complex (CPLX) overexpressing HEK293-EBNA cells, which do not assemble fibrillin-1 fibers, shows deposition of His_6_-tagged BMP-7 CPLX onto fibrillin-1 positive fibers. Scale bar: 50 µm. (B) Primary skin fibroblasts derived from *Fbn1* GT8 mice show impaired fibrillin-1 fiber assembly. Scale bar: 100 µm. (C) Co-culture of primary fibroblasts from *Fbn1* GT8 mice with HEK293-EBNA cells that overexpress BMP-7 CPLX. At equal cell numbers (1:1 ratio) between BMP-7 CPLX producing and wt fibroblasts no BMP-7 was detected in the cell culture supernatant as assessed by dot blot analysis indicating full sequestration of secreted BMP-7 CPLX under this condition. SDS-PAGE followed by western blot analysis of lysates of the corresponding cell layer showed no intracellular BMP pSMAD 1/5 signals indicating no induction of any BMP response. At equal ratios between *Fbn1^GT8/GT8^* fibroblasts and BMP-7 CPLX producing HEK293-EBNA cells (1:1 ratio), unsequestered BMP-7 CPLX was detected in the conditioned media inducing a pSMAD 1/5 response in the cell layer. (C) Increased BMP activity was measured in conditioned media from 3D-organotypic co-culture of *Fbn1* GT8 fibroblasts and wt keratinocytes via the *Id3* response in C2C12 cells. Increased presence of unsequestered BMP-7 protein was detected in conditioned media of *Fbn1* GT8 cultures by dot blot analysis after concentration of conditioned media over a heparin column. Error bars indicate mean ± SD from three independent experiments. ****P<0.0001. (D) Fibrillin-1 fiber degradation seen in Fig. 1B coincides with presence of increased pSMAD 1/5/8 signals in the dermis (blue arrowheads point to pSMAD 1/5/8 positive nuclei visualized by immunohistochemistry). Scale bar: 50 µm. (F) SDS-PAGE of skin lysates followed by western blot analysis showed increased pSMAD 1/5/8 signals at P7 and P14 in skin of *Fbn1* GT8 mice when fibrillin-1 fibers are degraded. Analyses of skin lysates from representative littermates are shown.

Dot blot analysis of cell culture supernatants showed that at equal cell numbers of wild-type fibroblasts and BMP-7 producing cells all produced BMP-7 CPLX was targeted to matrix fibers (Fig. 5C). However, when *Fbn1^GT8/GT8^*fibroblasts were used, unbound BMP-7 complex was present which induced a BMP response in the cell layer as indicated by increased signals of phospho-SMAD1/5 that are known signaling intermediates of BMP signaling (Figure 5C).

In a second experiment we analyzed BMP activity in the supernatant of a 3D organotypic co-culture system where fibroblasts from mutant mice were co-cultured with wild-type keratinocytes. By analysis of the cell culture supernatant we detected increasing amounts of endogenous non-sequestered BMP-7 protein and increased BMP activity as consequence of the increased presence of mutant *Fbn1* alleles (Fig. 5C).

In skin, we found at early postnatal time points when fibrillin-1 fibers are intact no indication of increased BMP signaling, however with the onset of fibrillin-1 fiber fragmentation increased BMP signaling indicated by pSMAD1/5/8 signals in the dermis and hair follicles was present (Fig. 5D, top). Further, western blot analysis of GT8 skin also showed evidence of increased BMP signaling indicated by pSMAD1/5/8 signals at P7 and P14 when degradation of fibrillin-1 was observed (Fig. 5D, bottom).

### Aberrant BMP signaling in *Fbn1^GT8/GT8^* aorta correlates with the onset of structural alterations in fibrillin-1 and collagen networks

Further we investigated thoracic aortas of *Fbn1^GT8/GT8^*mice for increased BMP signaling at postnatal time points when degradation of fibrillin fibers and structural alterations in the collagen network was observed (Fig. 1B, Fig. 3). At P7 *Fbn1^GT8/GT8^*aortic arches showed increased BMP signaling indicated by pSMAD1/5 staining with the strongest signals observed in the aortic valve region (Fig. 6A). *Fbn1^GT8/GT8^* mice at the age of P14 also showed increased pSMAD1/5 signals detected by western blot analysis (Fig. 6B) as well as mRNA upregulation of the BMP response gene *Id3* (Fig. 6C). However, qPCR analysis of TGF-β elements within the first two weeks of postnatal life did not indicate any increase of TGF-β signaling in the aorta of GT8 mice (Fig. S2).

**Figure 6:**
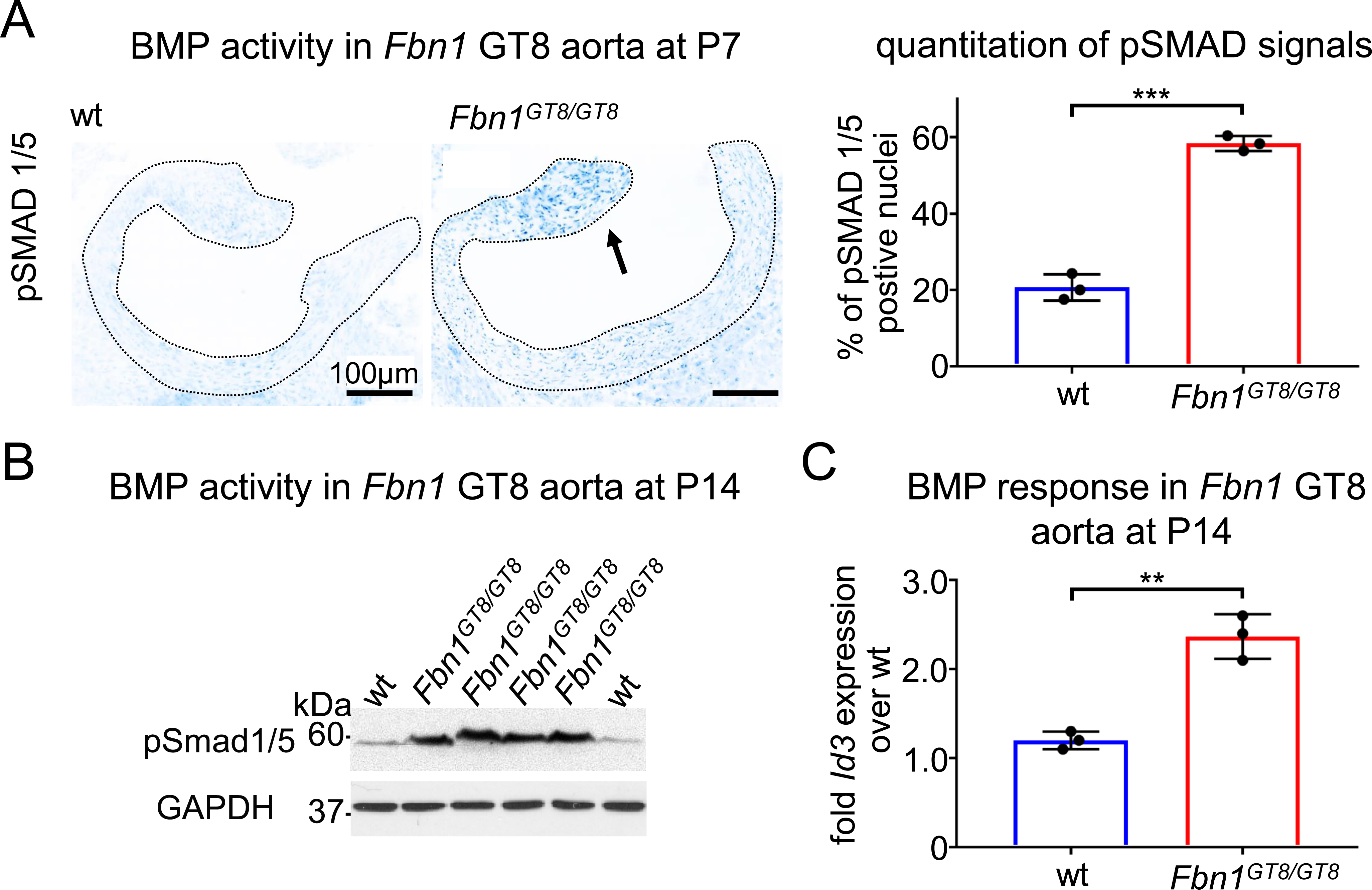
Aberrant BMP signaling in *Fbn1^GT8/GT8^* aorta correlates with the onset of structural alterations in fibrillin-1 and collagen networks. (A) (left) Immunohistochemistry signal in thoracic aorta sections at P7 employing a specific antibody against the BMP signaling intermediate pSMAD 1/5. For better visualization, the pSMAD 1/5 contrast was inverted by using ImageJ. *Fbn1^GT8/GT8^*mice showed the highest BMP activity in the aortic valve region (indicated by arrow). (right) For quantitation of pSMAD 1/5 signals, positively stained nuclei in wild-type (wt) and *Fbn1^GT8/GT8^* aortas were counted and divided by the total number of counted nuclei. (B) SDS-PAGE of lysates of *Fbn1^GT8/GT8^*aortic arches followed by western blot analysis employing an antibody against pSMAD 1/5 shows elevated BMP activity at P14. (C) Quantitative real-time PCR showed increased transcript levels of the BMP responsive gene *Id3* in aortic arches of *Fbn1^GT8/GT8^* mice at P14. The results are presented as mean ± SD from three mice per genotype. **P<0.001.

### MMP-13 is significantly increased in *Fbn1^GT8/GT8^* aortas

We recently reported that BMPs can induce MMP expression in HEK-293 cells and fibroblasts (18). To investigate whether the observed increased BMP activity would be causative for an increased expression of MMPs in the GT8 aorta we isolated primary vascular smooth muscle cells at P14 and analyzed transcript levels of various MMPs by qPCR in P14 aorta of *Fbn1^GT8/GT8^* mice. This analysis showed that *Mmp13* mRNA production at P14 was significantly upregulated (Fig. 7A). Further, analysis of *Mmp13* transcript levels of aortic arches isolated from *Fbn1^GT8/GT8^* mice showed that elevated mRNA expression was already detected at P7, but was significant at P14 (Fig. 7B). A similar induction of *Mmp13* transcript levels was also achieved when wild-type VSMCs isolated from aortic arches at P14 were stimulated with BMP-7, which is known to be expressed in human ascending aortic aneurysms (GEO Profile IDs: 44142292, 44142293, 44140680, 44140681) (Fig. 7C). Also, western blot analysis of lysates from dissected *Fbn1^GT8/GT8^* aortic arches and immunofluorescence analysis of thoracic aortic sections showed increased MMP-13 signals (Fig. 7D,E).

**Figure 7:**
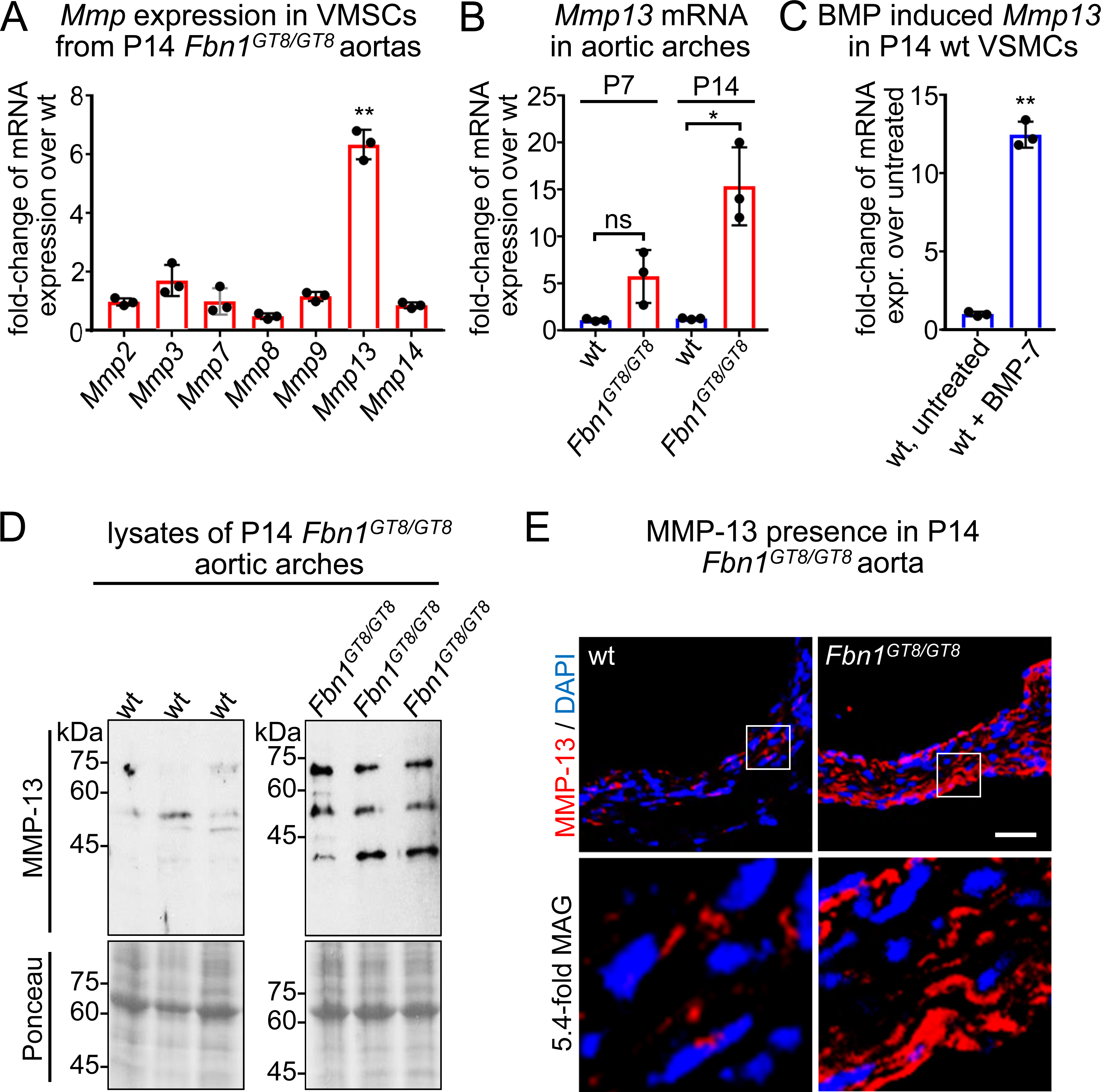
MMP-13 is significantly increased in *Fbn1^GT8/GT8^* aortas. (A) mRNA expression of various MMPs in VSMCs isolated from aortic arches of P14 wild-type (wt) and *Fbn1^GT8/GT8^* mice. **P<0.01. (B) Quantitative real-time PCR revealed increased *Mmp13* transcript levels at P14 in *Fbn1^GT8/GT8^*thoracic aorta. Results are presented as mean ± SD from three mice per genotype. Ns: not significant, *P<0.05. (C) BMP-7 stimulation of VSMCs isolated from P14 wt aortic induced *Mmp13* mRNA expression over untreated control. **P<0.001. (D) Western blot analysis using a specific antibody against MMP-13 showed elevated MMP-13 signals in lysates of P14 *Fbn1^GT8/GT8^* thoracic aortas. (E) Immunofluorescence analysis indicated a strong MMP-13 signal in *Fbn1^GT8/GT8^* thoracic aortas at P14 compared to wt littermate controls.

### Genetic ablation of *Mmp13* prevents aortic aneurysm formation in *Fbn1^GT8/GT8^* mice

To study an involvement of MMP-13 in aortic aneurysm formation in GT8 Marfan mice, the mutant *Fbn1* GT8 allele was bred onto a *Mmp13* null background. Dissected aortas showed a significant amelioration of aortic aneurysm growth in *Fbn1^GT8/GT8^*;*Mmp13^-/-^* compared to *Fbn1^GT8/GT8^* mice (Fig. 8A). At P11, *Fbn1^GT8/GT8^* mice showed an increased diameter in aortic annulus, sinuses of valsalva, and aorta ascendens in comparison to wild-type littermates as assessed by echocardiography (Fig. 8B).

**Figure 8:**
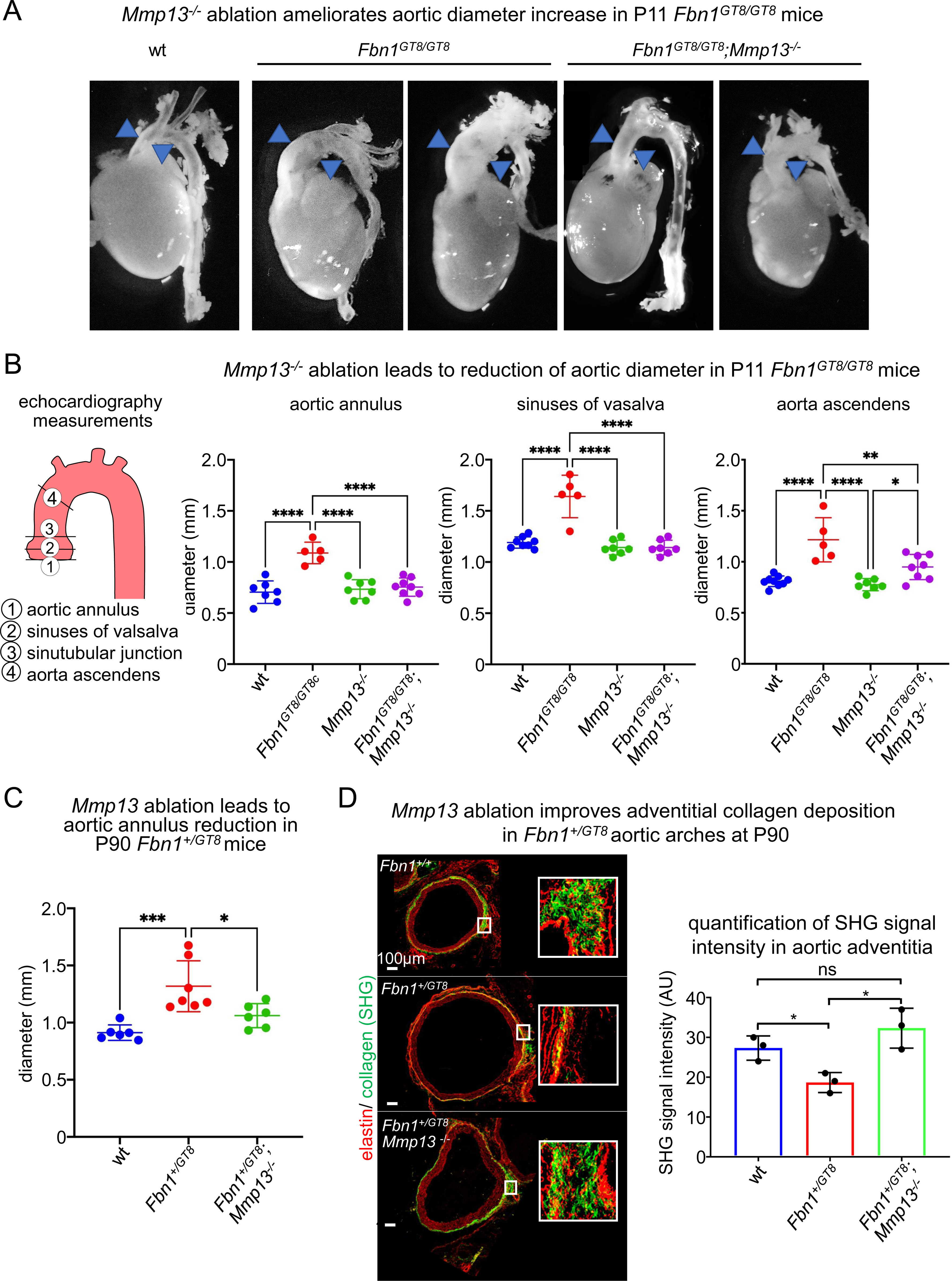
Genetic ablation of *Mmp13* prevents aortic aneurysm formation in *Fbn1^GT8/GT8^* mice. (A) Photographs of isolated P11 ascending aortas from wild-type (wt), *Mmp13^-/-^*, *Fbn1^GT8/GT8^,* and *Fbn1^GT8/GT8^*;*Mmp13^-/-^*mice showed that *Mmp13* ablation stops aortic diameter growth caused by the *Fbn1* GT8 allele. (B) (left) Scheme depicting the anatomical regions of the thoracic aorta (1–3) where diameter measurements were performed. (right) Aortic diameter measurements of P11 ascending aortas by echocardiography showed that *Mmp13* ablation leads to reduction of aortic diameter growth in *Fbn1^GT8/GT8^*mice. Each data point represents one analyzed animal. (C) *Mmp13* ablation leads to aortic annulus reduction in P90 *Fbn1^+/GT8^* mice shown as assessed by echocardiography measurements. (D) *Mmp13* ablation restores collagen presence in aortic adventitia in *Fbn1^+/GT8^* mice at three month of age. Thoracic aortas were harvested from 3 months old mice, paraffin-embedded, and sections were subjected to simultaneous SHG and immunofluorescence microscopy using an antibody against tropoelastin. The sections were scanned and the Maximum Intensity Projection (MIP) of each sample was considered for comparison. Signal quantification was performed using ImageJ. Results are presented as mean ± SD from three mice per genotype. Ns: not significant, *P<0.05.

A significant reduction of aortic annulus diameter was also found in *Fbn1^+/GT8^*;*Mmp13^-/-^*mice at P90 as indicated by echocardiography measurements (Fig. 8C). SHG analysis showed that adventitial collagen was restored in *Fbn1^+/GT8^*;*Mmp13^-/-^* aortas at P90 (Fig. 8D).

### *Mmp13* ablation restores collagen crosslink defects and decorin in *Fbn1* GT8 aorta

To gain further mechanistic insight into how mutant GT8-fibrillin-1 affects collagen integrity we subjected aortic arches from P90 *Fbn1^+/GT8^* mice to collagen crosslink analysis. Attempts to perform this analysis at earlier postnatal time points had failed due to the insufficient weight and therefore protein content of dissected thoracic aortas. Our analysis revealed a significant reduction of immature difunctional DHLNL (dihydroxylysinonorleucine) crosslinks in *Fbn1^+/GT8^* aortas that was normalized upon *Mmp13* ablation (Fig. 9A). Interestingly, collagen content as indicated by hydroxyproline content was normal in *Fbn1^+/GT8^* aortas and was not significantly changed upon *Mmp13* ablation (Fig. 9B). To investigate whether a reduction in collagen crosslinks leads to an increased collagen extractability and susceptibility to MMP-13 degradation, we isolated aortic arches aortas from *Fbn1^GT8/GT8^* mice at P10 and incubated them in MMP-13 reaction buffer in presence and absence of recombinant MMP-13 enzyme followed by western blot analysis (Fig. 9C). In MMP-13 reaction buffer containing 0.05% Triton X-100 more collagen I was extractable from *Fbn1^GT8/^*^GT8^ aortas compared to wild-type littermate controls (Fig. 9D). Extracted collagen I from *Fbn1^GT8/GT8^*aortas was also more susceptible to MMP-13 degradation (Fig. 9D).

**Figure 9:**
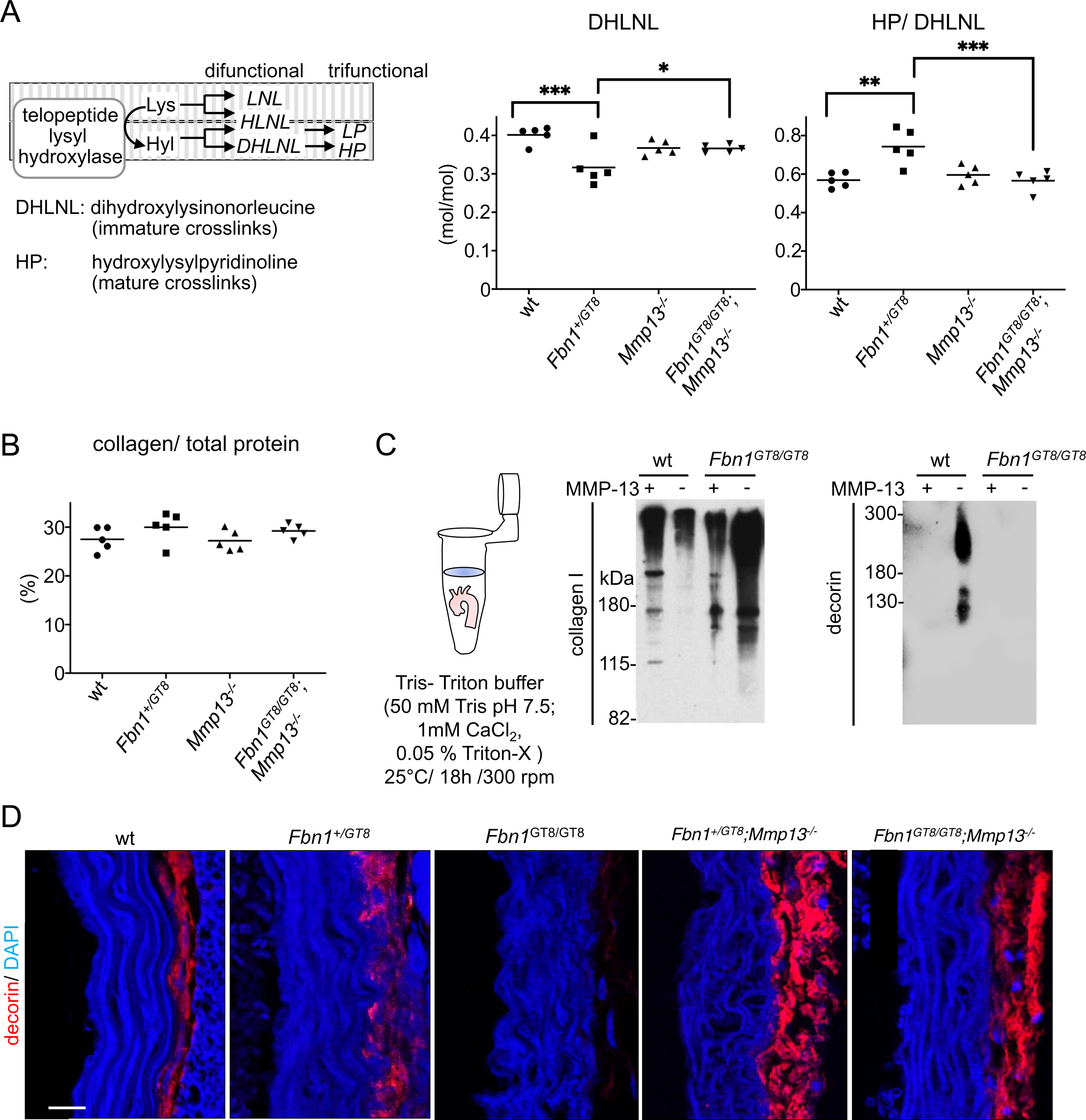
*Mmp13* ablation restores collagen crosslink defects and decorin in *Fbn1* GT8 aorta. (A) (left) Scheme illustrating mechanistic sequence for the establishment of difunctional and trifunctional collagen crosslinks. (right) Collagen crosslink analysis at P90 aortic arches of Fbn1+/GT8 mice showed are significant reduction in immature dihydroxylysinonorleucine (DHLNL) that was normalized upon *Mmp13* ablation. Each data point represents one animal. *P<0.05, **P<0.01, ***P<0.001. (B) Hydroxyproline analysis of P90 aortic arches from wild-type (wt), *Mmp13^-/-^*, *Fbn1^GT8/GT8^,* and *Fbn1^GT8/GT8^*;*Mmp13^-/-^*mice showed normal levels of collagen protein. (C) *In vitro* digestion assays of isolated aortic arches with recombinant MMP-13 followed by non-reducing SDS-PAGE and western blot analysis of collagen I and decorin. Collagen I was more extractable from *Fbn1^GT8/GT8^* aortas already in digestion buffer in absence of MMP-13 suggesting a reduced anchorage. Decorin was extractable in MMP-13 digestion buffer from wt but not *Fbn1^GT8/GT8^* aortas. Decorin extracted from wt aortas was quantitatively degraded upon MMP-13 addition. (D) Confocal immunofluorescence microscopy showed that decorin signal intensity is strongly diminished in the aortic adventitia of *Fbn1^+/GT8^*, and *Fbn1^GT8/GT8^* mice, but was restored upon *Mmp13* ablation. Scale bar: 50 µm.

Due to its prominent role in collagen alignment (34, 35) and therefore in aortic aneurysm formation (36) we also assessed the status of the small heparan sulfate proteoglycan (HSPG) decorin in *Fbn1* GT8 aortas. Decorin was extractable in wild-type aortas but quantitatively degraded upon MMP-13 addition (Fig. 9). However, in extracts of *Fbn1^GT8/GT8^* aortas no decorin was detectable suggesting its MMP-13 mediated degradation. To investigate whether *Mmp13* addition would restore decorin in *Fbn1* GT8 aortas, we employed immunofluorescence analysis on sections of dissected aortic arches. While increased presence of the *Fbn1* GT8 allele leads to a reduction of decorin signals in aortic adventitia, additional ablation of *Mmp13* restored decorin levels *Fbn1^+/GT8^* and *Fbn1^GT8/GT8^* aortas.

### Pharmacological inhibition of MMP-13 prevents aortic root growth in *Fbn1^+/GT8^* mice

To examine whether MMP-13 would could serve as potential therapeutic target in aortic aneurysm formation in Marfan syndrome, we treated *Fbn1^+/GT8^* at P30 for one month with a previously developed specific MMP-13 inhibitor with an IC_50_ of 1 nM that was determined in *in vitro* cleavage assays (). Pretests determined the most effective dose to be at 1mg/kg/day (Fig. S3). Evaluation of aortic diameters by echocardiography showed that inhibition of MMP-13 significantly prevented aortic annulus growth in *Fbn1^+/GT8^* mice similar to its genetic ablation in *Fbn1^+/GT8^*;*Mmp13^-/-^* mice (Fig. 10A).

**Figure 10:**
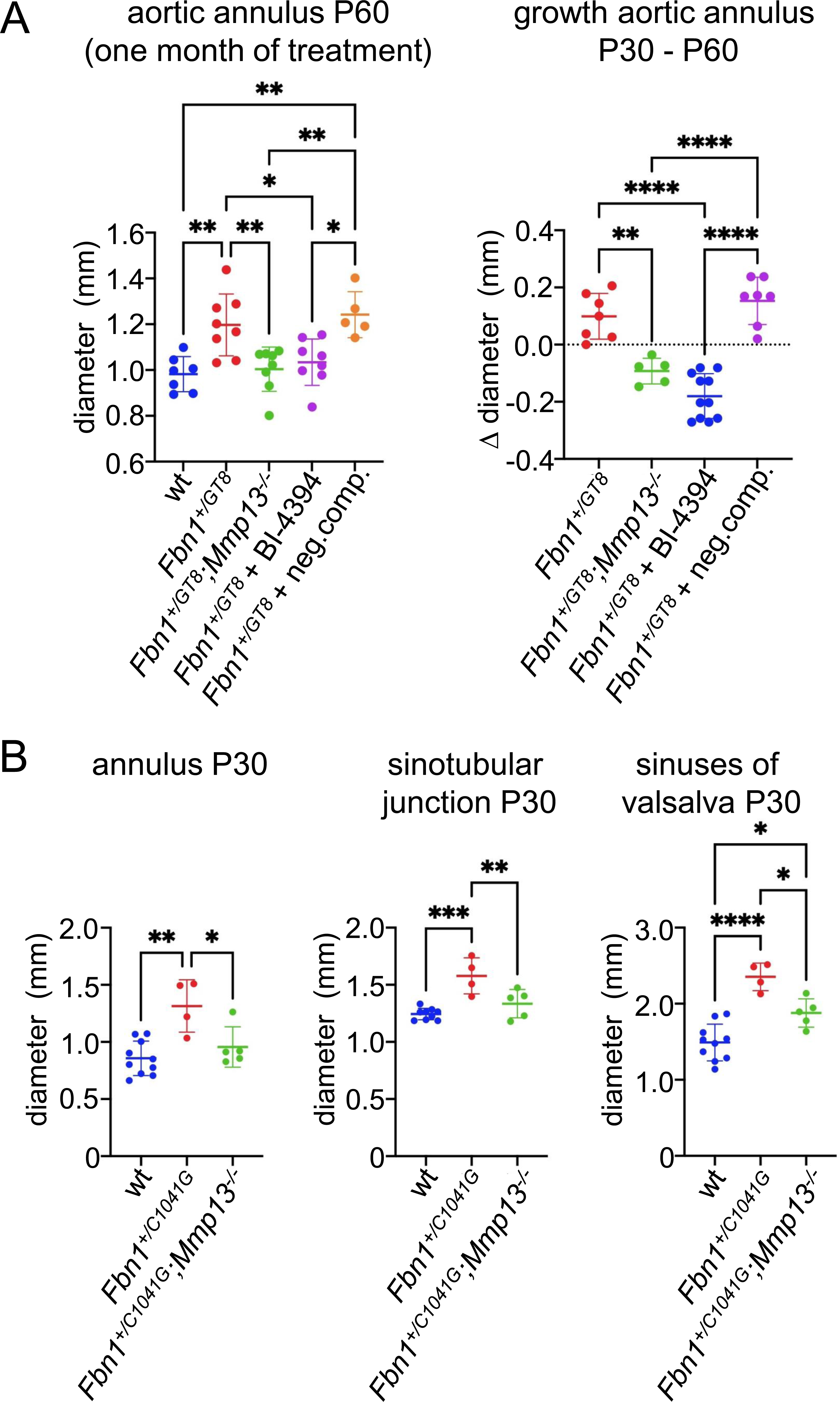
Evaluation of MMP-13 as therapeutic target in *Fbn1* GT8 and *Fbn1* C1049G Marfan mice. (A) Pharmacological inhibition of MMP-13 by application of BI-4394 (IC_50_=1nM) via osmotic minipumps at 1mg/kg/day from P30 until P60 showed significant amelioration of aortic annulus growth in *Fbn1^+/GT8^* mice similar to its genetic ablation in *Fbn1^+/GT8^*;*Mmp13^-/-^* mice. Each data point represents one animal. *P<0.05, **P<0.01, ****P<0.0001. (B) Genetic ablation of *Mmp13* in *Fbn1^+/C1041G^* mice caused significant amelioration of aortic diameter increase at aortic annulus, sinotubular junction, and sinuses of valsalva as assessed by echocardiography at P30. Each data point represents one animal. *P<0.05, **P<0.01, **P<0.001, ****P<0.0001.

### *Mmp13* ablation as promising strategy to reduce aneurysm growth in the *Fbn1^+/C1041G^* mouse model of *Fbn1* haploinsufficiency

To explore MMP-13 as a target also in other mouse models of MFS we bred the *Fbn1* C1041G allele onto a *Mmp13* null background. Echocardiography measurements at P30 indicated a significant reduction of aortic diameter increase in aortic annulus, sinuses of valsalva, and aorta ascendens in comparison to wild-type littermates (Fig. 10B). This suggests a general role of MMP-13 in the underlying pathological mechanism of aortic aneurysm formation caused by fibrillin-1 deficiency.

## Discussion

This work provided the first evidence that the collagenase MMP-13 plays a role during aneurysm formation in mouse models of MFS. Ablation of *Mmp13* in the dominant negative *Fbn1* GT8 mouse model of MFS resulted in significantly decreased diameter of the ascending aorta, and improved collagen fiber integrity. Also, *Mmp13* ablation in the haploinsufficient *Fbn1* C1041G mouse model led to a significant attenuation of aortic aneurysm growth suggesting MMP-13 as promising target in MFS. Moreover, pharmacological inhibition of MMP-13 showed a strong reduction in aortic aneurysm growth in *Fbn1* GT8 mice thereby proving its promoting role in aneurysm progression. Overall, our findings open up new therapeutic avenues to counteract aneurysm progression in MFS patients by application of already available drugs (19, 37, 38).

Our investigations allowed to gain new mechanistic insight into ECM-derived pathological mechanisms leading to the destruction of aortic wall integrity and aneurysm growth. FMF are known to be important scaffolds in the extracellular microenvironment of aortic resident cells as they not only define the biomechanical properties such as elasticity but also integrate the bioavailability of growth factors. Our gained results suggested a new working model of how fibrillin microfibril deficiency triggers degradative processes in aortic ECM architecture (Fig. 11). Structurally intact FMF sufficiently target and sequester BMP PD-GF CPLXs via specific PD-fibrillin-1 interactions, however, in the state of fibrillin-1 deficiency failed sequestration of BMP CPLXs leads to aberrant levels of bioactive BMP CPLXs that induce the expression of MMP-13. MMP-13 is known to degrade fibrillin-1 (39), decorin (40), and collagen I and III (41, 42) and thereby causes severe damage on the aortic ECM architecture leading to aortic aneurysm growth and dissection (Fig. 11). At the same time FMF-bound BMPs are activated and released upon MMP-13 mediated PD cleavage (18), suggesting the existence of a positive feed forward mechanism of MMP expression and BMP activation (2).

**Figure 11:**
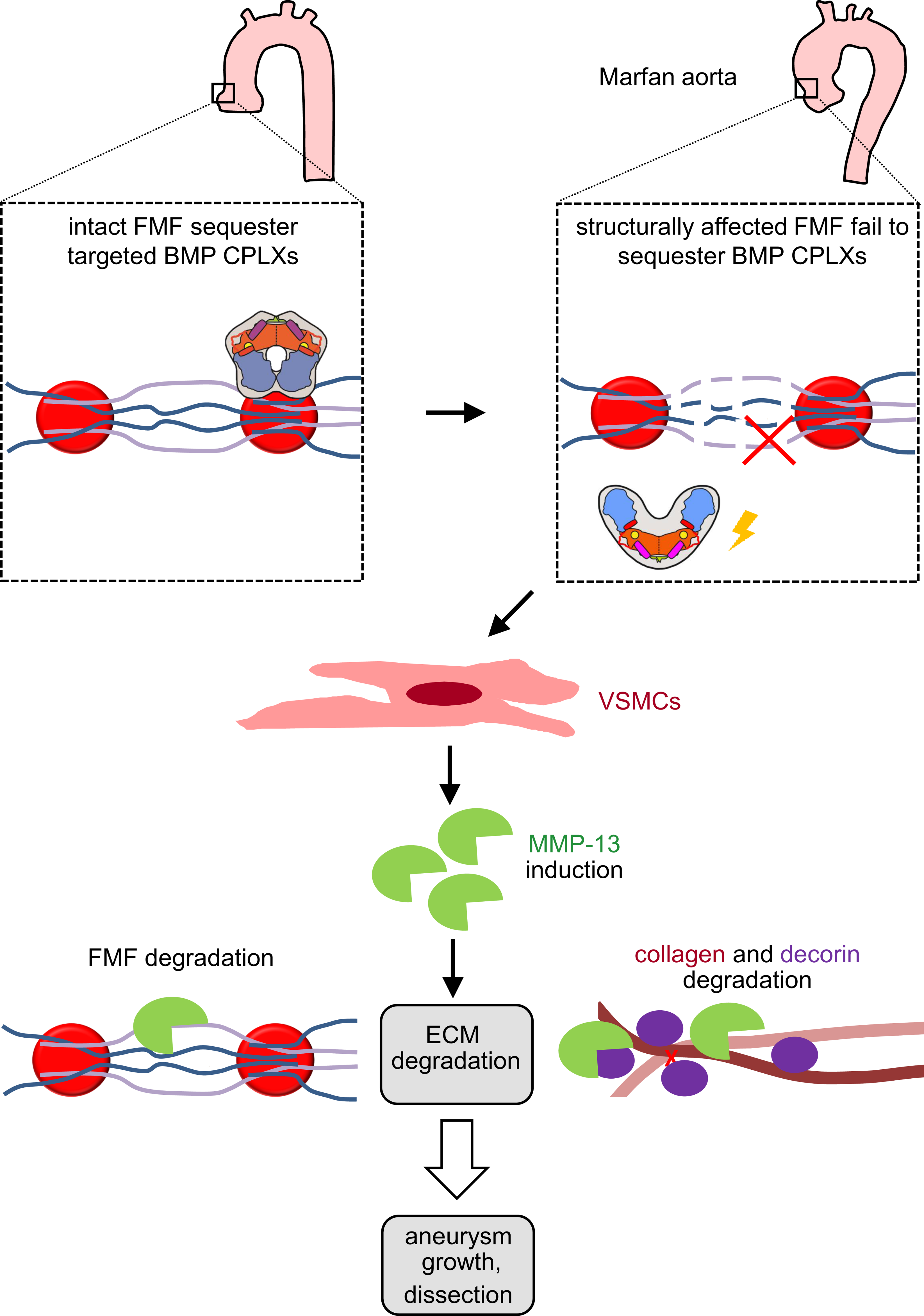
New model of how fibrillin microfibril deficiency triggers degradative processes in aortic ECM architecture. Structurally intact fibrillin microfibrils (FMF) target and sequester BMP prodomain-growth factor complexes (prodomain: blue, growth factor: orange). In the state of fibrillin-1 deficiency, failed sequestration of BMP CPLXs leads to aberrant levels of bioactive BMP CPLXs that induce the expression of MMP-13 which are known to not only degrade FMF but also decorin and the collagen network.

Our observations in *Fbn1^GT8/GT8^* mice suggest that in the first week of postnatal life (P0-P7), when fibrillin-1 fibers appear to be normal (Fig. 1B), fibrillin-2 compensates for the lack of intact fibrillin-1. However, between P7 and P14 fibrillin-2 production is turned off and GT8 fibrillin-1 is not able to maintain normal microfibril assembly. It has been hypothesized that fibrillins-1 and -2 functionally compensate for each other during embryonic development and that bundles of FMF represent heterogenic networks composed of both fibrillins (4, 43). To test this idea *Fbn1^-/-^*;*Fbn2^-/-^* mice were generated and the phenotypic outcomes compared the respective single knock-outs (22). Although, *Fbn1*^-/-^ tissues showed the presence of assembled FMF exclusively consisting of fibrillin-2, *Fbn1^-/-^*animals died within the perinatal period (22). Also, *Fbn1^-/-^;Fbn2^-/-^*mice are not viable, most likely due to a delay in elastic fiber formation in the ascending aorta during embryogenesis (22). Similarly, the importance of fibrillin-2 in *Fbn1* GT8 mice during fetal development was demonstrated, since *Fbn1^+/GT8^*, and *Fbn1^GT8/GT8^*mice cannot survive on a *Fbn2*^-/-^ background (5). These results indicate that during embryogenesis, fibrillin-1 and -2 have overlapping functions and that embryonic survival is fibrillin dosage-dependent. During fetal development, the expression of fibrillin-2 in bovine aorta was shown to be threefold higher than fibrillin-1 expression (44), however, during the perinatal period fibrillin-1 expression overtakes fibrillin-2 expression as shown in the mouse aorta (23), thereby substituting fibrillin-2 in the process of FMF assembly. Nevertheless, *FBN2* mutations lead to the autosomal dominant disorder, congenital contractual arachnodactyly (CCA, MIM#121050), characterized by a Marfanoid habitus such as tall stature, disproportional long limbs (dolichostenomelia) and also cardiovascular involvement such as thoracic aortic aneurysm formation.

Our analyses revealed a co-localization of collagen and elastin within the adventitia of wild-type aortas that was lost in the presence of mutant GT8-fibrillin (Fig. 3B). This indicates that proper alignment of both networks is required to provide sufficient structural integrity to the aortic wall. Already previously, similar profound defects in the collagen microarchitecture were found in thoracic aortic aneurysm (TAA) samples from MFS patients (31). Thereby, the collagen network in the aortic media showed only minor changes while a strongly disturbed collagen architecture was detected in the adventitial layer characterized by complete absence of normal collagen fibril organization and deposition of thin parallel collagen fibrils (31). Our co-immunolabeling analyses revealed a co-localization of fibrillin-1 and collagen III within the aortic adventitia and a partial co-distribution within the aortic media (Fig. 4D). This may imply that assembly of a proper FMF network is important for proper alignment of collagen fibers in the aortic adventitia and media.

MMP-13 belongs to the collagenase subgroup of the MMP family, and was previously found to be upregulated in abdominal aortic aneurysms (AAA) (45–47) and TAA (48). Our results suggest for the first time an involvement of MMP-13 in the initial steps of aortic aneurysm formation in MFS. Until now, only increased MMP-2 and MMP-9 activity was measured in aortic extracts of MFS mouse models at adult age (49–51), however, conflicting results were obtained in MFS patients. For instance, increased MMP-2 activity was found in some MFS patients (52) while in others reduced activity levels of MMP-2 were measured (53). Until now, aneurysm formation is believed to occur through elastic fiber break down as a first step, followed by collagen degradation. However, our investigations in the *Fbn1^GT8/GT8^*aorta at early postnatal time points suggest that collagen and elastic fiber degradation occur simultaneously (Fig. 2,3). MMP-13 is known to not only degrade components of the aortic collagen network such as collagen I, III, and decorin, but also crucial proteins of the FMF/elastic fiber network such as fibronectin (54), fibrillin-1 (39), and perlecan (55). All of these proteins have a vital contribution to aortic wall stability and are implicated in aortic aneurysm formation and dissection (56, 57).

In this context, decorin is an interesting MMP-13 substrate as it plays an important functional role in the aortic ECM. Decorin was shown to be reduced in abdominal aortic aneurysms (AAA) resulting in vessel wall instability thereby predisposing the vessel to rupture (36). Also, treatment with recombinant decorin attenuated the formation and rupture of Ang II-induced AAA in mice by reinforcing the aortic wall (36). It is known that interactions between the core protein of decorin and collagen govern the rate and extent of collagen fibril growth thereby modulates their tensile strength (34, 35). Thereby any alterations of decorin abundance or distribution would be expected to influence the pattern of fibril formation and, thus, alignment of crosslinking sites (35). Our results showed that MMP-13 mediated reduction of intact decorin correlates with a reduced presence of DHLNL crosslinks (Fig. 9). This suggests that in decorin is required to maintain proximity to functionally reactive collagen crosslinking residues in the *Fbn1* GT8 aorta. Further, since decorin also interacts with FMF (58, 59), it may function as bridge molecule to mediate the functional interdependence of collagen and elastin fiber networks. This identified interdependence is likely relevant to human pathology since also previous reports described a functional link between fibrillin-1 and decorin deficiency in fibroblast cultures from patients with neonatal MFS (60, 61).

Aortic dissection is exclusively attributed to collagen degradation, as impaired elastic fibers alone were not sufficient to induce aortic dissection in *ex vivo* models (62). Increased turnover of type III collagen has been reported in aortic aneurysms in patients with AAA, and monitored by the small peptide PIIINP which is liberated into extracellular fluids (63). Furthermore, it has been suggested that imbalances between collagen degradation and synthesis significantly weakens ECM network integrity leading to aortic wall rupture (64). Therefore, our results are in line with these previous findings, suggesting that reduced collagen integrity is the most important driver of aneurysm growth in TAA.

The stimulation of VSMCs with BMP-7 GF led to increased MMP-13 expression (Fig. 7C), suggesting that increased levels of MMP-13 protein and mRNA in the GT8 aorta are due to aberrant activation of BMP signaling. Also, our BMP stimulation experiments with HEK293 and primary murine fibroblasts showed a BMP induced upregulation of MMP-13 transcript levels (18). BMP-dependent induction of MMP-13 expression was also found to be implicated in ECM degradation during developmental or connective tissue disease processes. For instance, BMP-2 and -4 stimulation of primary human fibroblast caused upregulation of MMP-1, -2, -3, and -13 which was proposed to be a mechanism in melanoma invasion (65). During chondrogenesis, BMPs regulate terminal differentiation during which hypertrophic chondrocytes remove the collagen matrix via the upregulation of MMP-13 (66). In osteoarthritis (OA), elevated BMP levels in damaged cartilage promote cartilage degeneration by stimulating MMP-13 expression (67).

MMP-13 is not produced in most adult human tissues, but plays a prominent role in ECM remodeling during fetal development (68). In zebrafish, for instance, *mmp13* morpholino knockdown affected heart formation (69). In humans, endothelial cells from aortic and pulmonary valves were found to overexpress MMP-13 for tissue maturation (70). Overexpressed MMP-13 was also found in myxomatous mitral valve disease (71), a condition characterized by loss of collagen and glycosaminoglycan (GAG) accumulation (72). These findings point to a critical role of MMP-13 for heart integrity during embryogenesis, and in states of disease in postnatal life. Not much is known about the functional relationship between BMP signaling and MMP-13 activity in cardiac tissues. Abrogation of MMP-13 in the atherosclerosis ApoE null mouse model led to an improvement of collagen deposition within atherosclerotic plaques (73). Also, during embryogenesis both MMP-13 and BMP-2 were found to be specifically expressed in the cardiac valves, where BMP-2 was shown to modulate cell plasticity in valve precursor cells by stimulating the expression of aggrecan and Sox9 and MMP-13 modulates matrix remodeling (70, 74).

Previously, it was shown that aortas of adult *Fbn1^+/C1041G^*showed evidence of increased aberrant TGF-β activity (21). Interestingly, treatment of *Fbn1^+/C1041G^* mice at two month of age with TGF-β neutralizing antibodies or losartan, an angiotensin II type I (AT1) receptor blocker which lowers TGF-β activity, prevented and reversed aortic aneurysm formation (21). In *Fbn1^GT8/GT8^* mice we did not observe any signs of increased TGF-β at postnatal time points when ECM degradation and aortic aneurysms were already established. This suggests that dysregulated TGF-β activity is not involved in aneurysm establishment in *Fbn1^GT8/GT8^* mice. Our findings are in line with new hypothesis proposing that aberrantly increased TGF-β activity is a secondary consequence of major ECM remodeling events in the course of aortic aneurysm formation, which suggests that initiation of aortic disease in MFS precedes activation of TGF-β. This hypothesis was tested by breeding *Fbn1* C1041G mice to mice carrying a *Tgfbr2* knock-out allele under the control of a VSMC-specific Cre-driver. In this study the aorta of *Fbn1* C1041G mice at the age of two months developed aneurysms and showed no difference in read-outs for canonical and non-canonical TGF-β signaling pathways compared to wild-type controls, as previously stated (21). In addition, *Fbn1* C1039G mice on a *Tgfbr2* null background showed exacerbated aneurysm progression (75). These findings suggest that aneurysm formation in MFS is not primarily caused by increased TGF-β signaling.

Our results suggest that aberrant activation of BMP signaling (Fig. 6) initiates the onset of aneurysm formation. These findings are in line with our previous findings: (1) BMPs bind directly and specifically via their PDs to fibrillin-1 and -2 (7, 32), and may therefore serve as direct sensors for FMF integrity; (2) Non sequestered BMP PD-GF CPLXs are bioactive (76), but targeting to fibrillin-1 renders the inactive (33); (3) MMPs mediated activation of BMPs occurs via cleavage of the PD at specific sites (18). Therefore, our results may explain how dysregulated BMP signaling due to fibrillin-1 deficiency might be causative for aneurysm onset in *Fbn1* GT8 mice. However, treatment of MFS patients by BMP signaling antagonism may not be a promising strategy. It has been shown that BMP/TGF-β signaling is required for the maintenance of elastic fiber homeostasis (77). Therefore, it can be assumed that general antagonism of BMP signaling is likely to lead to aggravation of aneurysm formation.

Overall, our results suggest antagonism of MMP-13 as a promising strategy to specifically stop aortic root diameter increase in the Marfan aorta. Since MMP-13 represents an embryonic MMP that is not expressed under normal conditions throughout postnatal life, its inhibition promises to be safe with no unwanted side effects. Comprehensive studies subjecting other *Fbn1* mouse models of MFS will have to follow to fully explore MMP-13 antagonism in the pathology of aneurysm growth caused by fibrillin-1 deficiency.

## Nonstandard abbreviations

AAA: abdominal aortic aneurysm
BMP: bone morphogenetic protein
BM: basement membrane
CPLX: complex
EBNA: Epstein-Barr virus nuclear antigen
eGFP: enhanced green fluorescent protein
ELISA: enzyme linked immunosorbent assay
ECM: extracellular matrix
FMF: fibrillin microfibrils
GAG: glycosaminoglycans
GDF: growth and differentiation factor
GT8: green truncated from founder 8
GF: growth factor
HEK: human embryonic kidney
HSPG: heparan sulfate proteoglycan
MAB: monoclonal antibody
MFS: Marfan syndrome
MMP: matrix metalloproteinase
PCA: Principal component analysis
PD: prodomain
RT: room temperature
SHG: second harmonic generation
TAA: thoracic aortic aneurysm
TEM: transmission electron microscopy
TGF: transforming growth factor
TMB: tetramethylbenzidine

## Acknowledgments

We would like to thank Beatrix Martiny and Christian Jüngst from the CECAD Imaging Core Facility at the University of Cologne. Beatrix Martiny performed EM sample preparation, production of ultrathin sections, and recording of EM photographs. Christian Jüngst adjusted microscopy settings and assisted with recording of SHG signals. We further would like to thank Boehringer Ingelheim for providing the MMP-13 inhibitor BI-4394 and the negative compound BI-4395.

## Author contribution statement

L.A.Z. A.G.F. and G.S. conceived the study. L.A.Z., A.G.F., D.M., S.G., A.V.Z., G.P., N.P., T.v.B., D.S.B. J.M., K.S.L., J.B. performed research and analyzed data. P.Z. and M.G. provided essential reagents. M.G., P.Z., and S.B. provided expert advice. G.S. wrote and edited the manuscript. G.S. acquired funding for this study.

## Funding

This work was supported by the Deutsche Forschungsgemeinschaft (DFG) TRR259 (397484323) to G.S. (project B09), M.G. (project B08), and to S.B. (project A04). In addition, DFG funding was provided by SFB829 (73111208) to P.Z. (project B04) and G.S. (project B12). J.M. and K.S.L. acknowledge funding from the Ministry of Baden-Württemberg for Economic Affairs, Labor and Tourism (3-4332.62-NMI/65), and the DFG (INST 2388/33-1, INST 2388/64-1, and Germany’s Excellence Strategy-EXC 2180-390900677).

## Conflict of interest

The authors declare no conflict of interest.

## Notes

### Competing Interest Statement

The authors have declared no competing interest.

