## supplemental Fig1 and 2 for "Targeting of MMP-13 prevents aortic aneurysm formation in Marfan mice"

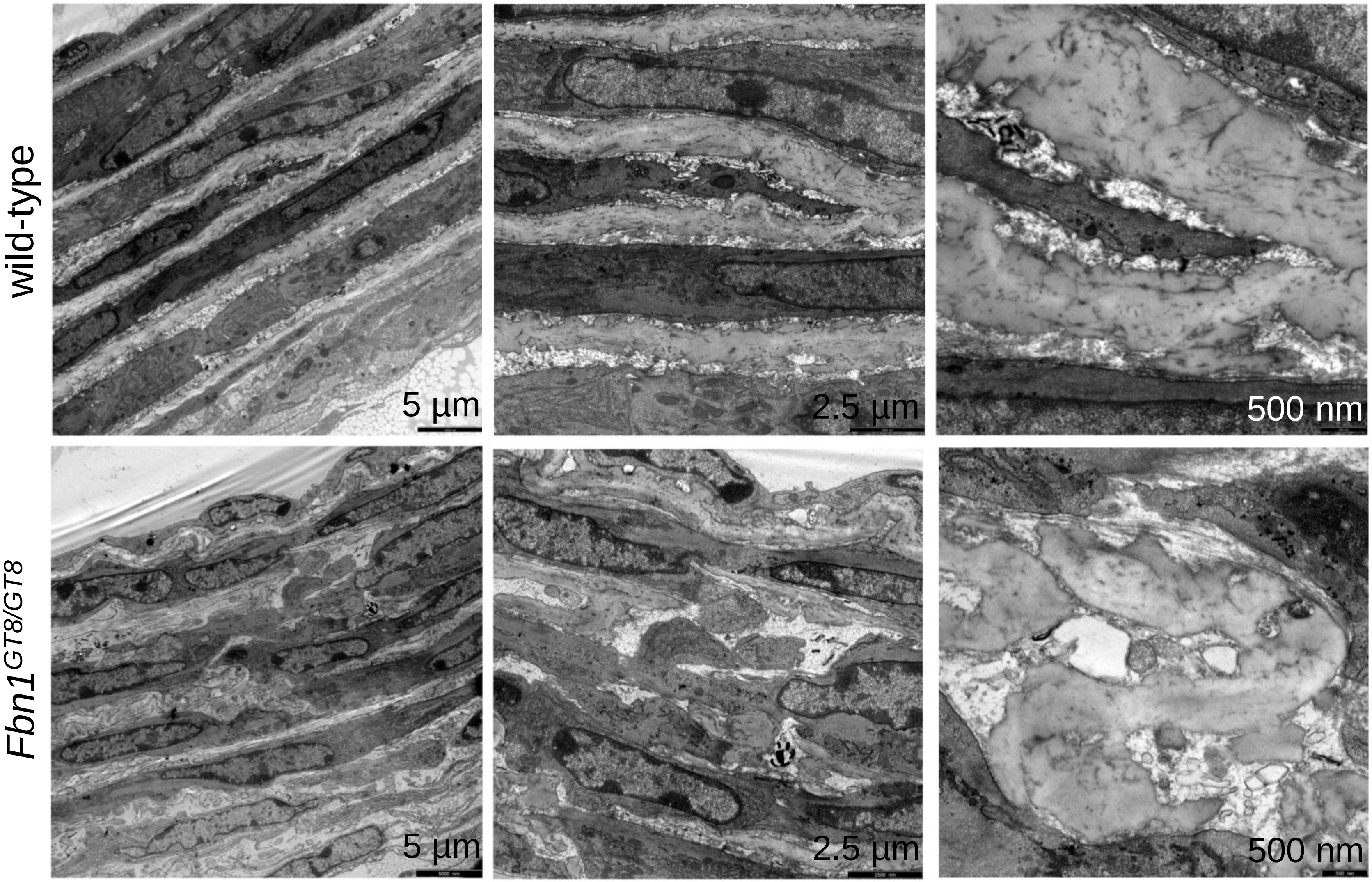


**Figure S1: Ultrastructural analysis of *Fbn1*$^{GT8/GT8}$ aortas at P9 reveals structural alterations of elastic lamellae but no changes in VSMC morphology.** Transmission electron microscopy of P9 aortic sections showing discontinous elastic lamellae and increased spacing of the intra lamellar space in *Fbn1*$^{GT8/GT8}$ aortas leading to a reduced physical contact of VSMCs with adjacent elastic lamellae. Closer inspection did not reveal any major morphological changes between VSMCs in *Fbn1*$^{GT8/GT8}$ and wild-type control aortas.

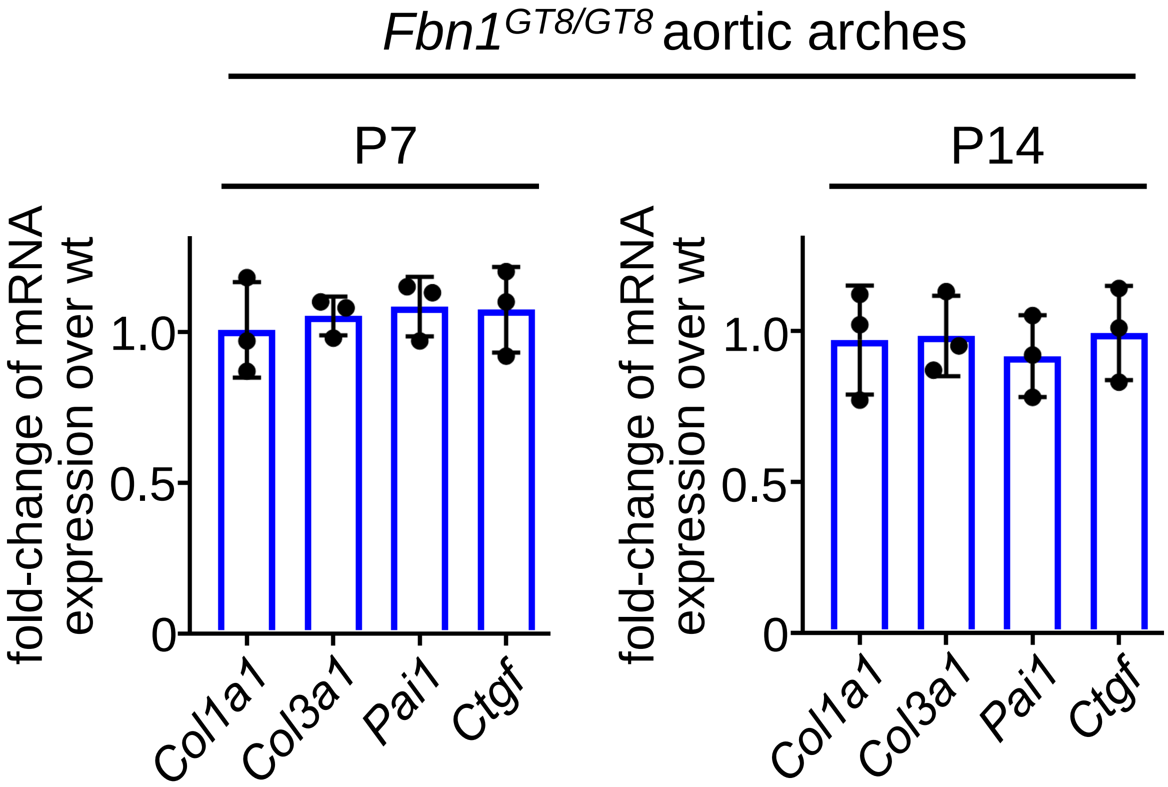


**Figure S2: Analysis of transcript levels of TGF-β read-out genes in *Fbn1*$^{GT8/GT8}$ aortic arches at P7 and P14 showed no significant differences.**
